# The anterior cingulate cortex and its role in controlling contextual fear memory to predatory threats

**DOI:** 10.1101/2021.02.02.429419

**Authors:** Miguel Antonio Xavier de Lima, Marcus Vinicius C. Baldo, Fernando A. Oliveira, Newton Sabino Canteras

## Abstract

Predator exposure is a life-threatening experience and elicits learned fear responses to the context in which the predator was encountered. The anterior cingulate area (ACA) occupies a pivotal position in a cortical network responsive to predatory threats, and it exerts a critical role in processing fear memory. Ours results revealed that the ACA is involved in both the acquisition and expression of contextual fear to predatory threat. Overall, the ACA can provide predictive relationships between the context and the predator threat and influences fear memory acquisition through projections to the basolateral amygdala and perirhinal region and the expression of contextual fear through projections to the dorsolateral periaqueductal gray. Our results expand previous studies based on classical fear conditioning and open interesting perspectives for understanding how the ACA is involved in processing contextual fear memory to ethologic threatening conditions that entrain specific medial hypothalamic fear circuits (i.e., predator- and conspecific-responsive circuits).

## INTRODUCTION

Predator exposure is a life-threatening experience and elicits innate fear behaviors as well as learned fear responses to the context in which the predator was encountered (Blanchard et al., 1989, 2001; Ribeiro-Barbosa et al., 2005). Recent studies revealed a cortical network that is responsive to predatory threats and exerts a critical role in processing fear memory. Thus, the caudal prelimbic area (PL), rostral part of the anterior cingulate area (ACA), medial visual area (VISm), and the ventral part of retrosplenial area (RSPv) form a highly interconnected circuit that presents a differential increase in Fos expression in response to predator exposure. Cytotoxic lesions of the elements of this cortical circuit apparently had no impact on innate fear responses during predator exposure but had a profound impact on contextual fear memory largely disrupting learned contextual fear responses in a predator-related environment (de Lima et al., 2019).

The ACA occupies a central position in this cortical network and establishes dense bidirectional connections with all other elements of the circuit. Importantly, lesions in the ACA had a larger impact on decreasing learned contextual fear responses versus lesions of the other elements of this cortical network (de Lima et al., 2019). However, at this point, it is not clear how the ACA is involved in processing predator-related fear memory.

Previous studies in the literature using fear conditioning to physically aversive stimuli (i.e., footshock) reported important roles for the ACA in fear memory. The ACA seems to be necessary for the acquisition of contextual fear. Pretraining inactivation of the ACA blocked fear acquisition (Tang et al., 2005; Bissière et al., 2008), and pretraining activation of the ACA using a mGluR agonist (Bissière et al., 2008) enhanced fear learning thus suggesting an involvement of the ACA in the acquisition of fear responses.

Of relevance, the insertion of an empty temporal gap, or trace, between the CS and UCS makes learning the association critically dependent on the prefrontal cortex. Recording studies in trace fear conditioning revealed units called “bridging cells” in the prefrontal cortex that maintain firing during the trace intervals perhaps reflecting the maintenance of attentional resources during the CS-USC interval (Gilmartin et al., 2014). These findings from trace fear conditioning suggest that the prefrontal cortex strengths the predictive value of the available cue to influence memory storage. The ACA is also required for memory consolidation. Infusion of the protein-synthesis inhibitor anisomysin in the ACA impairs memory consolidation of recent and remote memory (Einarsson and Nader, 2012), and virally mediated disruption of learning-induced dendritic spine growth in the ACA impairs memory consolidation (Vetere et al., 2011). In fact, the ACA has been largely associated with the encoding and retrieval of contextual fear memory at remote time points (Frankland et al., 2004; Kitamura et al., 2017; Abate et al., 2018).

Here, we examined how that ACA mediates predator fear memory. To this end, we exposed mice to cats and investigated innate and contextual fear responses. We started by asking whether the ACA is involved in the acquisition and/or expression predator fear memory and applied pharmacogenetic silencing in the ACA during exposure to the predator or to the context. Next, combining retrograde tracing and Fos protein immunostaining, we examined the pattern of activation of ACA source of inputs during acquisition and expression of predator fear responses. We could see how the ACA would be able to combine predator and contextual cues particularly during the acquisition phase. To complement these findings, during the cat exposure, we applied optogenetic inhibition to the anteromedial thalamic nucleus > ACA path, which putatively relays information related to the predator cues, and examined the effect on the acquisition of predator fear memory.

Next, using optogenetic silencing and functional tracing combining Fluoro Gold and Fos immunostaining, we examined how the ACA entrains selected targets to influence acquisition or expression of predator fear memory. The data help to clarify how the ACA influences both the acquisition and expresion of predator fear memory. Overall, the ACA offers predictive relationships between the threaten stimuli and the context to influence memory storage in amygdalar and hippocampal circuits. It also has a role in memory retrieval and the expression of contextual fear. Our results open interesting perspectives for understanding how the ACA is involved in processing contextual fear memory to predator threats as well as other ethologic threatening condition such as those seen in the confrontation with a conspecific aggressor during social disputes.

## RESULTS

### The ACA is involved in both the acquisition and expression of contextual fear to predator threat

To specifically evaluate the contribution of the ACA in the acquisition and expression of contextual fear to a predatory threat, we used designer receptors exclusively activated by designer drug (DREADD) to selectively silence the activity of the ACA during the cat exposure and during the exposure to the predatory context (Figure 1). We used adeno-associated virus (AAV) expressing Gi-coupled hM4Di fused with mCitrine fluorescent protein (AAV5-hSyn-HA-hM4D(Gi)-IRES-mCitrine). We bilaterally injected the viral vector (AAV5-hSyn-HA-hM4D(Gi)-IRES-mCitrine) or a vector expressing only the fluorescent protein for the control group (AAV5-hSyn-eGFP) into the ACA. Patch clamp experiments showed that transfected neurons (G_i_-DREADD neurons) hyperpolarized as they underwent extracellular CNO and presented a significant decrease in the triggering of action potentials after 10 μM CNO (S Fig. 3). Four weeks after the viral injection, the group of animals injected with virus expressing Gi-coupled hM4Di or the control virus received CNO intraperitoneally 30 min before the cat exposure or the exposure to the predatory context.

**Figure 1.**
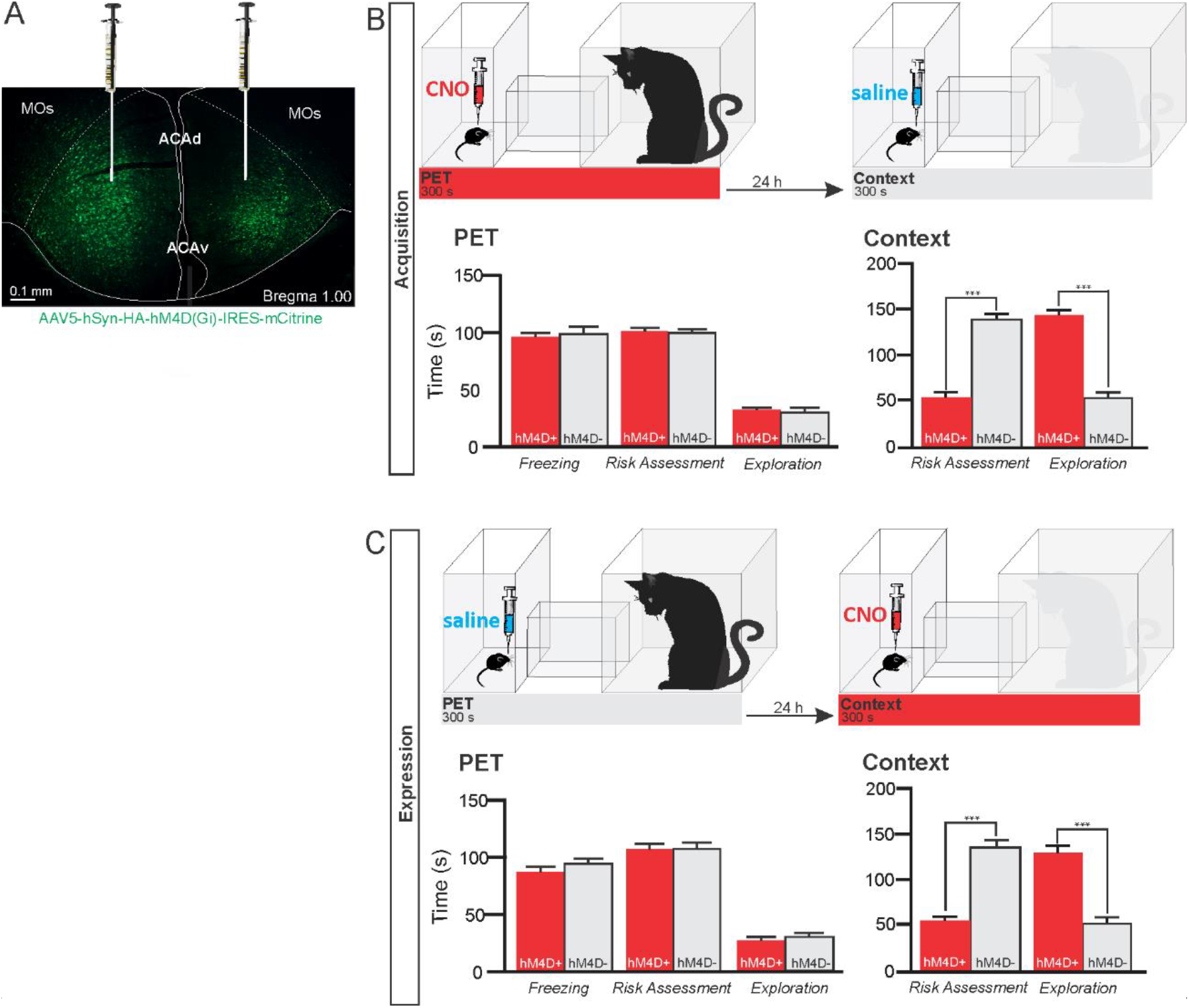
Pharmacogenetic inhibition of the ACA during acquisition and expression of contextual fear to predator threat. **A.** Fluorescence photomicrograph illustrating the bilateral injection in the ACA of a viral vector expressing inhibitory DREADD (hM4D Gi) fused with mCitrine. **B.** Pharmacogenetic inhibition of the ACA during the cat exposure (acquisition phase): (top) Experimental design, and (bottom) mean (± SEM) values of the behavioral responses during Predator Exposure (PET) and Predatory Context (Context). For inhibition during PET- Groups: hM4D+ (*n*=7) and hM4D-(*n*=6). **C.** Pharmacogenetic inhibition of the ACA during the predatory context (expression phase): (top) Experimental design, and (bottom) mean (± SEM) values of the behavioral responses during Predator Exposure (PET) and Predatory Context (Context). For inhibition during Context -Groups: hM4D+ (*n*=7) and hM4D-(*n*=5). Data are shown as mean ± SEM. 2 x 2 univariate ANOVAs for freezing and three-way ANOVAs for risk assessment and exploration followed by Tukey’s HSD test post hoc analysis (***p<0.001). Abbreviations: ACAd, anterior cingulate area, dorsal part; ACAv, anterior cingulate area, ventral part; CNO, clozapine N-oxide; MOs, secondary motor area; PET, predator exposure test.

The results showed that compared to the control group, animals expressing Gi-coupled hM4Di treated before the cat exposure did not change innate responses but significantly reduced contextual fear responses (Figure 1B). Thus, during exposure to the predatory context, the animals presented a significant decrease in the risk assessment responses and increase in fearless exploration (Figure 1B). Likewise, CNO treatment before the exposure to the predatory context also significantly reduced risk assessment and increased exploration in animals injected with virus expressing Gi-coupled hM4Di (Figure 1C). The results suggest that silencing the ACA does not influence innate fear responses but affects both the acquisition and the expression of contextual fear to predator threat. Complete statistical analysis for the behavioral data of ACA pharmacogenetic inhibition is found in the supplementary materials (S7a).

### Pattern of activation of the different sources of inputs to the ACA during the exposure to the cat and to the predatory context

Considering the influence of the ACA in the acquisition and expression of contextual fear to predatory threat, we next examined the pattern of activation of the different sources of inputs to the ACA during the exposure to the cat and to the predatory context. To this end, animals received unilateral deposits of a retrograde tracer (Fluoro Gold) in the ACA. One week later, groups of animals were exposed either to the cat only (Figure 2) or to the predatory context (S Fig. 5) and perfused 90 min later. For each one of these groups, we examined the percentage of Fluoro Gold labeled cells expressing Fos protein to see the activation pattern of the ACA inputs during the acquisition and expression of contextual fear. During the cat exposure, among the cortical inputs, the ventral retrosplenial (RSPv) and anteromedial visual (VISam) areas presented the largest percentage of FG retrogradely-labeled cells expressing Fos (around 50% of the retrogradely labeled cells); the other cortical inputs to the ACA, including the medial orbital (ORBm) and prelimbic (PL) areas, displayed close to 30% of double labeled cells (Figure 2C). According to our results, inputs to the ACA from the RSPv and VISm seem particularly activated during the acquisition phase of predator fear memory and are known to be involved in computing contextual landmarks (see Discussion).

**Figure 2.**
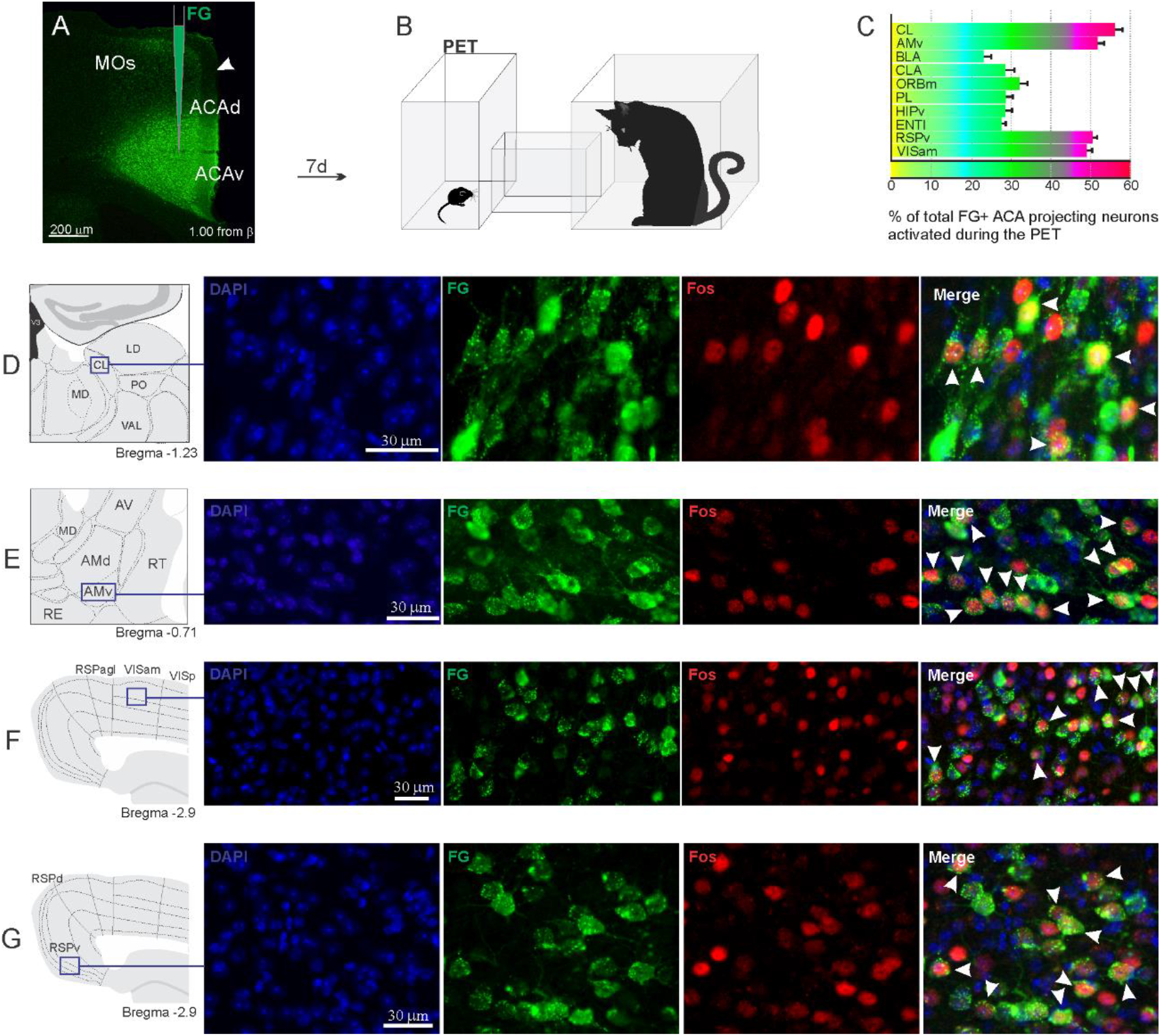
Pattern of activation of the different sources of inputs to the ACA during the exposure to the cat. Animals received unilateral deposit of a retrograde tracer (Fluoro Gold) in the ACA (n=6, **A**). Seven days later, animals were exposed to the cat (PET, **B**) and perfused 90 min after. **C.** Bar chart presents, for each designated structure, the proportion of activated neurons (*FG/Fos* double labeled cells) during the PET condition, among the total of FG retrogradely labeled cells (error bars indicate 95% confidence interval for a proportion). **D-G.** Schematic drawings from the *Allen Mouse Brain Atlas* to show the sites containing the largest proportion of *FG/Fos* double labeled cells, followed by fluorescence photomicrographs illustrating DAPI-staining, FG labeled cells in green (Alexa 488), Fos protein positive cells labeled in red (Alexa 594) and merged view of the FG and FOS labeled cells, where arrow heads indicate FG/FOS double labeled cells. Abbreviations: ACAd, anterior cingulate area, dorsal part; ACAv, anterior cingulate area, ventral part; AMd, anteromedial thalamic nucleus, dorsal part; AMv, anteromedial thalamic nucleus, ventral part; AV, anteroventral nucleus of thalamus; BLA, basolateral amygdalar nucleus; CL, central lateral nucleus of the thalamus; CLA, claustrum; ENTl, entorhinal area, lateral part; FG, Fluoro gold; HIPv, hippocampus, ventral part; LD, lateral dorsal nucleus of the thalamus; MD, mediodorsal nucleus of the thalamus; MOs, secondary motor area; ORBm, orbital area, medial part; PET, predator exposure test; PL, prelimbic area; PO, posterior complex of the thalamus; RE, nucleus of reuniens; RSPagl, retrosplenial area, lateral agranular part; RSPd, retrosplenial area, dorsal part; RSPv, retrosplenial area, ventral part; RT, reticular nucleus of the thalamus; VAL, ventral anterior-lateral complex of the thalamus; VISam, anteromedial visual area; VISp, primary visual area.

In addition, during cat exposure, a particularly large percentage of FG/Fos double labeled cells were found in the central lateral (CL) and ventral anteromedial thalamic (AMv) nuclei—both of these contained more than 50% retrogradely labeled expressing Fos protein (Figure 2C). Notably, the CL and AMv are likely to convey information regarding the predatory threat (see Discussion). We also found that, during cat exposure, inputs to the ACA from the basolateral amygalar nucleus, ventral hippocampus, lateral entorhinal area, and claustrum presented 25% to 30% FG/FOS double labeled cells (Figure 2C).

Exposure to the predatory context resulted in a different profile on the activation of the inputs to the ACA (S Fig. 5). Thus, during exposure to the predatory context, all cortical inputs to the ACA as well as the inputs from the basolateral amygdala and lateral entorhinal area presented close to 20% to 30% of FG-labeled cells expressing Fos protein (S Fig. 5C). In the thalamus, the AMv displayed a relatively low percentage of double labeled cells (close to 20%) whereas the central lateral nucleus contained close to 40% of the retrogradely-labeled cells expressing Fos protein (S Fig. 5C). Taken together, these findings revealed a differential activation among the inputs to the ACA during the acquisition and expression of the predator fear contextual memory suggesting the inputs from the VISm, RSPv, and AMv particularly activated during the acquisition phase whereas the CL inputs were largely activated during both the acquisition and expression of contextual fear memory to predatory threat.

### AM > ACA projection controls the acquisition of contextual fear responses to predator threats

Considering that the AMv is one of the inputs to the ACA presenting the largest activation during cat exposure, we employed projection-based silencing approach to investigate the effect of the inhibition of the AM > ACA projection on the acquisition of contextual fear to cat exposure. For the AM > ACA projection photoinhibition, adeno-associated viral (AAV) vectors encoding halorhodopsin-3.0 fused with mCherry fluorescence protein (AAV5-hSyn-eNpHR3-mCherry) or AAV control vectors not expressing halorhodopsin-3.0 encoding mCherry fluorescence protein (AAV5-hSyn-mCherry) were injected bilaterally into the AM (Figure 3 A, B). To test the efficiency of light-induced hyperpolarization in eNpHR3.0-expressing neurons, the patch clamp experiments in neurons transfected with halorhodopsin reveled a robust hyperpolarization with 585 nm light on with clear linear regression due to different light intensities (S Fig. 4).

**Figure 3.**
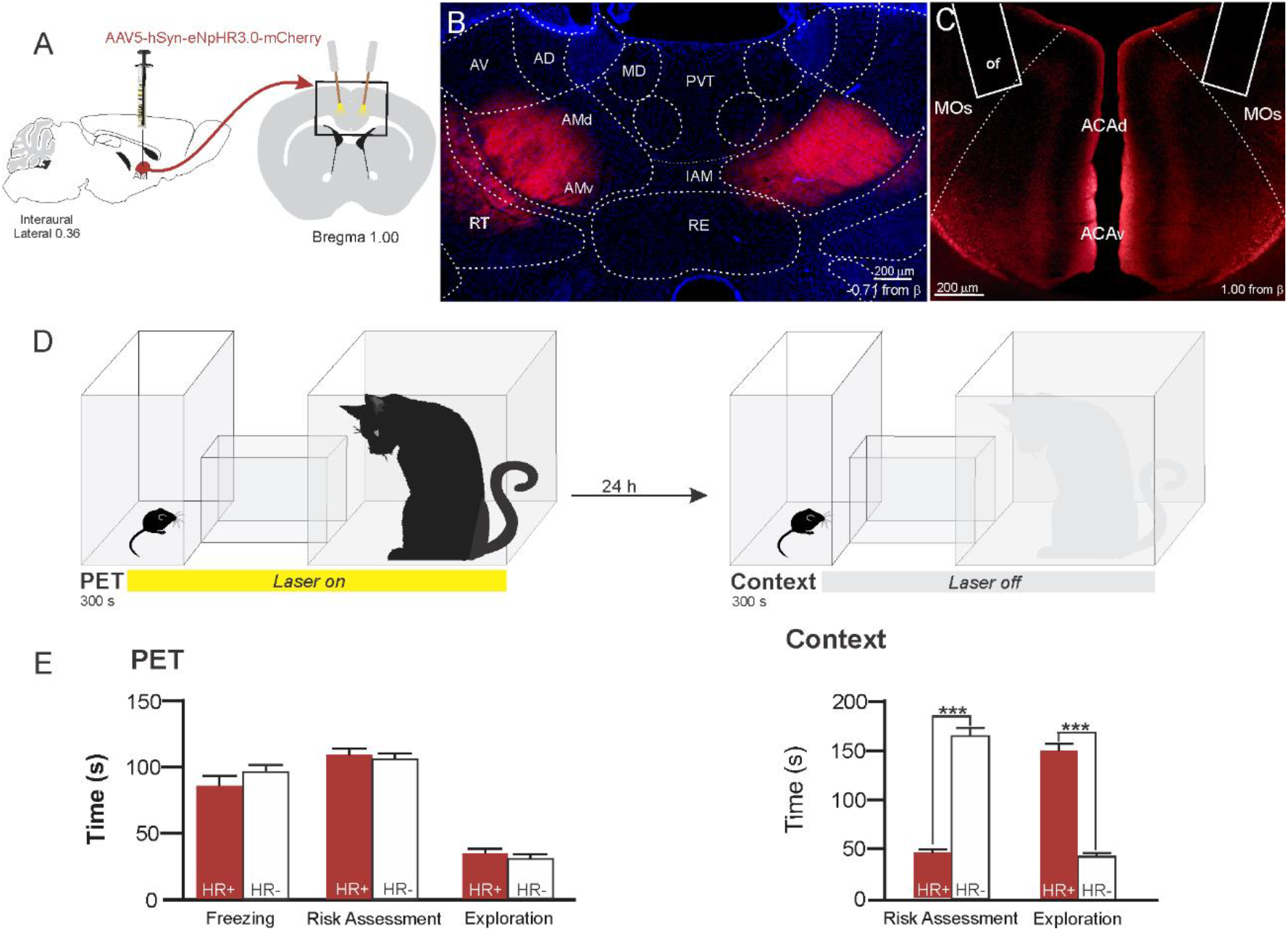
Optogenetic inhibition of anteromedial thalamic nucleus > ACA pathway during cat exposure. **A.** Schematics showing the location of the bilateral AAV viral vector injection in the anteromedial thalamic nucleus (AM) and the position of bilateral optical fibers implanted in the ACA. **B**. Fluorescence photomicrographs illustrating the bilateral injection in the AM of a viral vector expressing halorhodopsin-3.0 (eNpHR3.0) fused with mCherry. **C.** Fluorescence photomicrograph illustrating the mCherry anterograde labeled projection to the ACA (*of* - the optic fibers’ tips position). **D.** Experimental design. **E.** Mean (± SEM) values of the behavioral responses during Predator Exposure (PET) and Predatory Context (Context). Data are shown as mean ± SEM. Groups: HR+ (*n*=8) and HR-(*n*=7). One-way ANOVA for freezing and 2×2 ANOVAs for risk assessment and exploration followed by Tukey’s HSD test post hoc analysis (***p<0.001). Abbreviations: ACAd, anterior cingulate area, dorsal part; ACAv, anterior cingulate area, ventral part; AD, anterodorsal nucleus of thalamus; AMd, anteromedial thalamic nucleus, dorsal part; AMv, anteromedial thalamic nucleus, ventral part; AV, anteroventral nucleus of thalamus; IAM, interanteromedial nucleus of the thalamus; MD, mediodorsal nucleus of the thalamus; MOs, secondary motor area; of, optical fiber; PET, predator exposure test; PVT, paraventricular; nucleus of the thalamus; RE, nucleus of reuniens; RT, reticular nucleus of the thalamus.

A 589 nm laser light was continually delivered to the ACA during the 5 min of cat exposure through surgically implanted dual-fiber optic elements (Figure 3C). Specific viral expression in the AM was confirmed for each mouse after behavioral testing by microscopic observation. Our results showed that photoinhibition of the AM > ACA projection during cat exposure did not change innate fear responses but significantly reduced contextual fear response and impaired the acquisition of contextual fear response to a predator threat (Figure 3E). Thus, the animals presented a significant decrease in the risk assessment responses and increase in fearless exploration during exposure to the predatory context (Figure 3E). Complete statistical analysis for the behavioral data of the photoinhibition of the AM > ACA projection is supplied in the supplementary material (S7b). Here, the AM may convey predator cues to the ACA processed in the hypothalamic predator-responsive circuit; this finding nicely suits the known roles of the ACA in attention and assessment of cue salience during the learning processes (see Discussion).

### Differential roles of ACA projection targets on the acquisition and expression of contextual fear to predator threat

This work used optogenetic silencing and functional tracing combining Fluoro Gold and Fos immunostaining. We examined how the ACA entrains selective targets to influence acquisition or expression of predator fear memory. Among the ACA targets, we start by exploring projections to the basolateral amygdala and the perirhinal area—these are key targets for the prefrontal cortex to influence associative plasticity and memory storage in the amygdala and hippocampus (see Gilmartin et al., 2014). Next, we examined the ACA projection to the post-subiculum (POST), which has a large influence on the medial performant path, and conceivably in the memory processing (Ding, 2013). Finally, we explored the projection to the dorsolateral periaqueductal because studies have shown its putative roles both in the acquisition and the expression of contextual fear memory to predator threat (Cezário et al., 2008; de Andrade Rufino et al., 2019); therefore, a likely ACA target to influence such responses. As illustrated in the Supplementary Figure 6, the pattern of ACA projection to each one of these selected targets were examined using viral tracing.

We employed projection-based silencing of the ACA projections to each one of the selected targets and tested the effect of photoinhibition during the acquisition phase and the expression of contextual fear to animals previously exposed to the cat. To this end, adeno-associated viral (AAV) vectors encoding halorhodopsin-3.0 fused with mCherry fluorescence protein (AAV5-hSyn-eNpHR3-mCherry) or AAV control vectors not expressing halorhodopsin-3.0 encoding mCherry fluorescence protein (AAV5-hSyn-mCherry) were injected bilaterally into the ACA (Figs 4A, 5A, 6A, and 7A). A 589 nm laser light was continually delivered through surgically implanted dual-fiber optic elements to each one of the ACA selected targets (Figs 4B, 5B, 6B and 7B) during the 5 min exposure to the cat or the predatory context.

**Figure 4.**
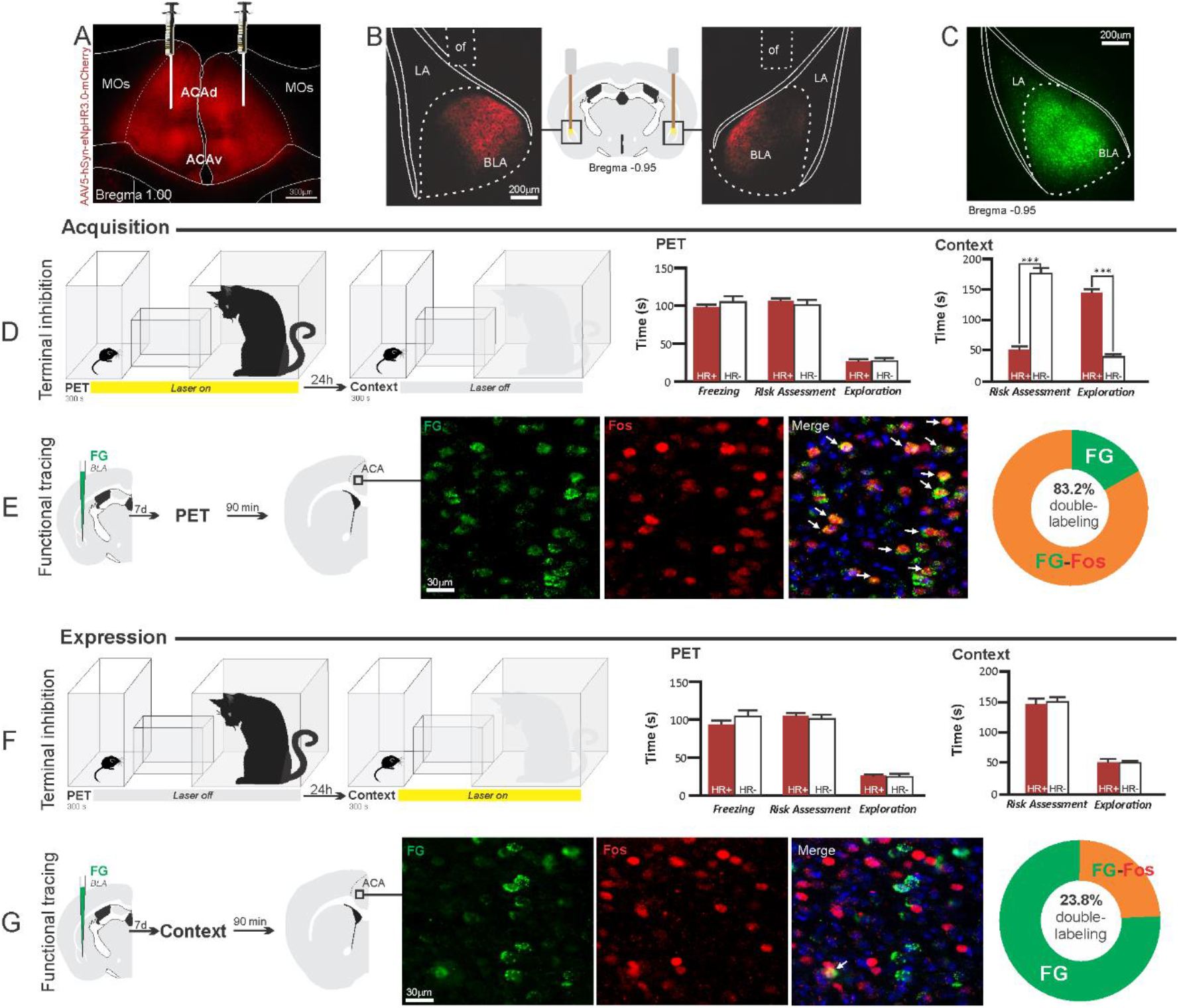
Optogenetic silencing and functional tracing of the ACA > BLA pathway during the acquisition and expression of contextual fear to predator threat. **A.** Fluorescence photomicrograph illustrating the bilateral injection in the ACA of a viral vector expressing halorhodopsin-3.0 (eNpHR3.0) fused with mCherry. **B.** Schematic drawing (center) and fluorescence photomicrographs showing the ACA projections to the BLA and location of bilateral optical fibers implanted close to the basolateral amygdala (*of* - optic fibers’ tips position). **C.** Fluorescence photomicrograph illustrating the FG injection in the BLA for the functional tracing. **D and F.** Optogenetic silencing of the ACA > BLA pathway during the cat exposure (**D**) or predatory context (**F**). Experimental design (on the left) and mean (± SEM) values of the behavioral responses during Predator Exposure (PET) and Predatory Context (on the right). For silencing during PET condition - Groups: HR+ (*n* = 8) and HR-(*n* = 6); for silencing during Context condition - Groups: HR+ (*n* = 8) and HR-(*n* = 6). Data are shown as mean ± SEM. 2 x 2 ANOVA for freezing and three-way ANOVAs for risk assessment and exploration followed by Tukey’s HSD test post hoc analysis (***p<0.001). **E and G.** Functional tracing of the ACA > BLA pathway during the cat exposure (acquisition phase, **E**) and the predatory context (expression phase, **G**). Left - Experimental design: unilateral FG injection in the BLA, and 7 days later perfusion 90 min after the cat exposure (n=4, **E**) or the context exposure (n=4, **G**). Center - Fluorescence photomicrographs illustrating, in the ACA, FG labeled cells in green (Alexa 488), Fos protein positive cells labeled in red (Alexa 594) and merged view of the FG and FOS labeled cells (arrows indicate FG/FOS double labeled cells). Right – Graphic representation of the percentage of FG/Fos double labeled cells in the ACA. Abbreviations - ACAd, anterior cingulate area, dorsal part; ACAv, anterior cingulate area, ventral part; BLA, basolateral amygdalar nucleus; FG, Fluoro gold; LA, lateral amygdalar nucleus; MOs, secondary motor area; of, optical fiber; PET, predator exposure test.

**Figure 5.**
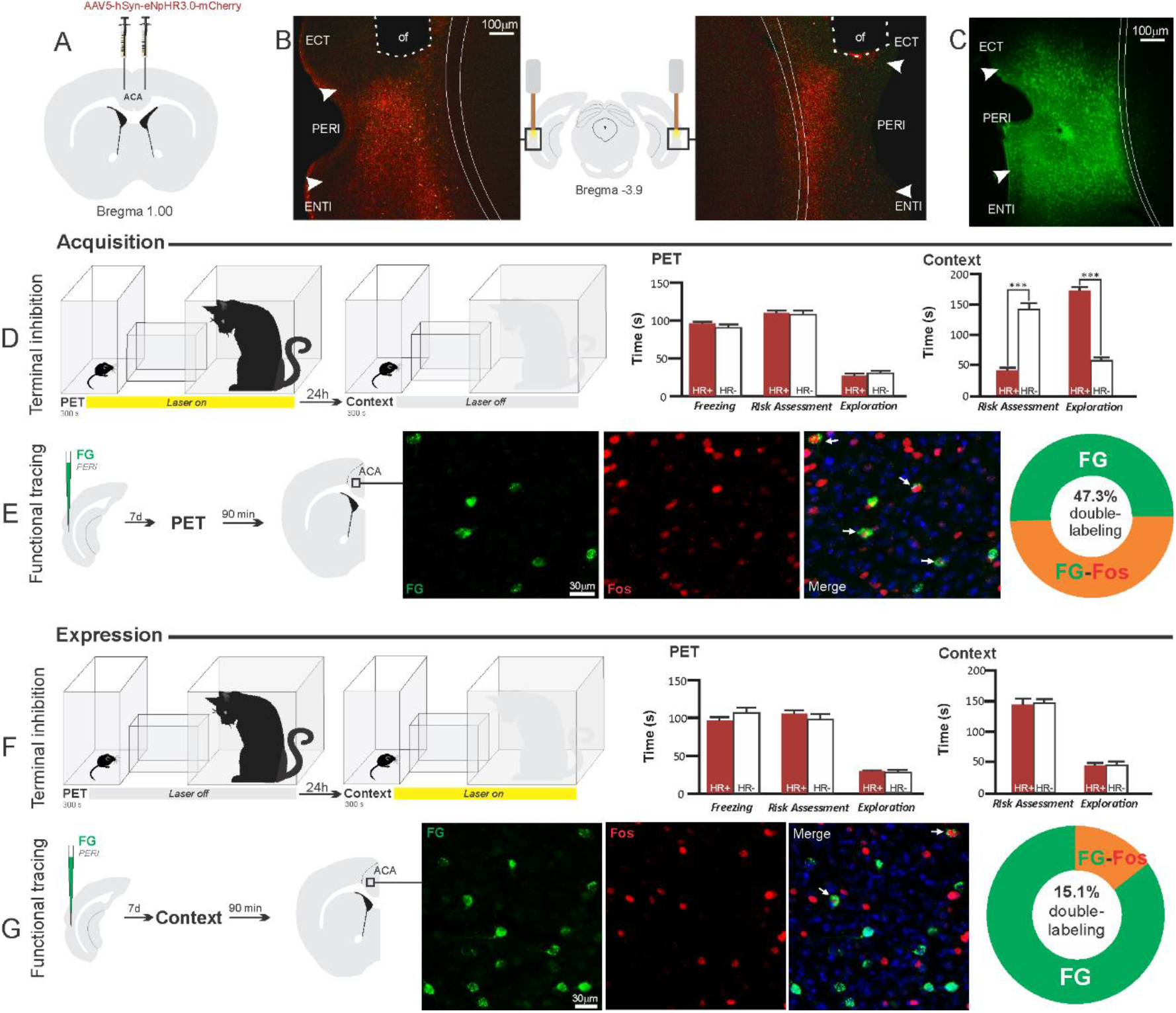
Optogenetic silencing and functional tracing of the ACA > PERI pathway during the acquisition and expression of contextual fear to predator threat. **A.** Schematic drawing illustrating the bilateral injection in the ACA of a viral vector expressing halorhodopsin-3.0 (eNpHR3.0) fused with mCherry. **B.** Schematic drawing (center) and fluorescence photomicrographs showing the ACA projections to the PERI and location of bilateral optical fibers implanted close to the PERI (*of* - optic fibers’ tips position). **C.** Fluorescence photomicrograph illustrating the FG injection in the PERI for the functional tracing. **D and F.** Optogenetic silencing of the ACA > PERI pathway during the cat exposure (**D**) or predatory context (**F**). Experimental design (on the left), and mean (± SEM) values of the behavioral responses during Predator Exposure (PET) and Predatory Context (on the right). For silencing during PET condition - Groups: HR+ (*n*=6) and HR-(*n*=5); for silencing during Context condition - Groups: HR+ (*n*=6) and HR-(*n*=5). Data are shown as mean ± SEM. 2 x 2 ANOVA for freezing and three-way ANOVAs for risk assessment and exploration followed by Tukey’s HSD test post hoc analysis (*p<0.001). **E and G.** Functional tracing of the ACA > PERI pathway during the cat exposure (acquisition phase, **E**) and the predatory context (expression phase, **G**). Left - Experimental design: unilateral FG injection in the PERI, and 7 days later perfusion 90 min after the cat exposure (n=4, **E**) or the context exposure (n=4, **G**). Center – Fluorescence photomicrographs illustrating, in the ACA, FG labeled cells in green (Alexa 488), Fos protein positive cells labeled in red (Alexa 594) and merged view of the FG and FOS labeled cells (arrows indicate FG/FOS double labeled cells). Right – Graphic representation of the percentage of FG/Fos double labeled cells in the ACA. Abbreviations - ACA, anterior cingulate area; ECT, ectorhinal area; FG, Fluoro gold; of, optical fiber; PET, predator exposure test; ENTl, entorhinal area, lateral part; PERI, perirhinal area.

**Figure 6.**
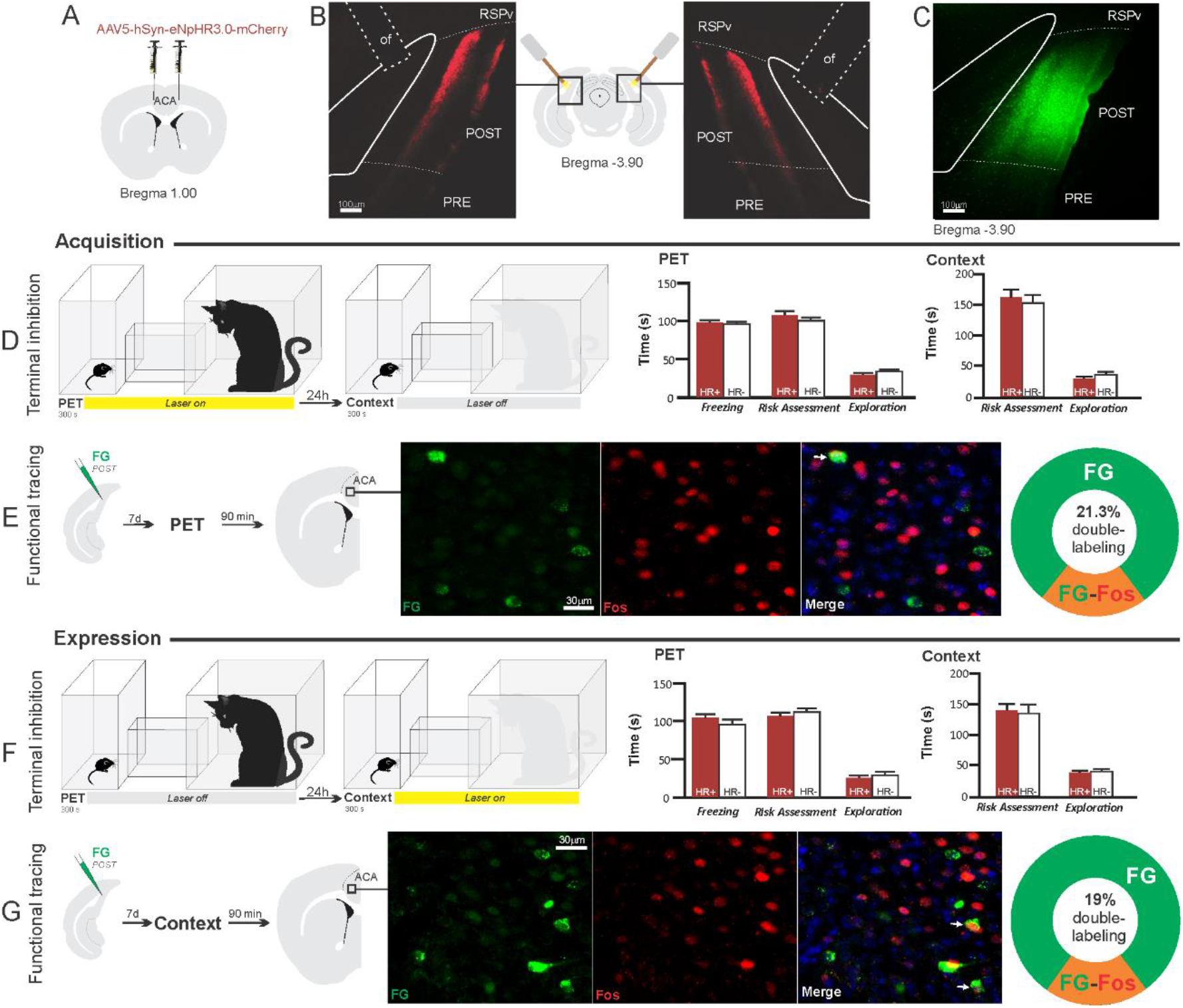
Optogenetic silencing and functional tracing of the ACA > POST pathway during the acquisition and expression of contextual fear to predator threat. **A.** Schematic drawing illustrating the bilateral injection in the ACA of a viral vector expressing halorhodopsin-3.0 (eNpHR3.0) fused with mCherry. **B.** Schematic drawing (center) and fluorescence photomicrographs showing the ACA projections to the POST and location of bilateral optical fibers implanted close to the POST (*of* - optic fibers’ tips position). **C.** Fluorescence photomicrograph illustrating the FG injection in the POST for the functional tracing. **D and F.** Optogenetic silencing of the ACA > POST pathway during the cat exposure (**D**) or predatory context (**F**). Experimental design (on the left) and mean (± SEM) values of the behavioral responses during Predator Exposure (PET) and Predatory Context (on the right). For silencing during PET condition - Groups: HR+ (*n*=6) and HR-(*n*=5); for silencing during Context condition - Groups: HR+ (*n*=7) and HR-(*n*=5). Data are shown as mean ± SEM. 2 x 2 ANOVA for freezing and three-way ANOVAs for risk assessment and exploration followed by Tukey’s HSD test post hoc analysis. **E and G.** Functional tracing of the ACA > POST pathway during the cat exposure (acquisition phase, **E**) and the predatory context (expression phase, **G**). Left - Experimental design: unilateral FG injection in the POST, and 7 days later perfusion 90 min after the cat exposure (n=4, **E**) or the context exposure (n=4, **G**). Center - Fluorescence photomicrographs illustrating, in the ACA, FG labeled cells in green (Alexa 488), Fos protein positive cells labeled in red (Alexa 594) and merged view of the FG and FOS labeled cells (arrows indicate FG/FOS double labeled cells). Right – Graphic representation of the percentage of FG/Fos double labeled cells in the ACA. Abbreviations - ACA, anterior cingulate area; FG, Fluoro gold; of, optical fiber; PET, predator exposure test; POST, postsubiculum; PRE, presubiculum; RSPv, retrosplenial area, ventral part.

**Figure 7.**
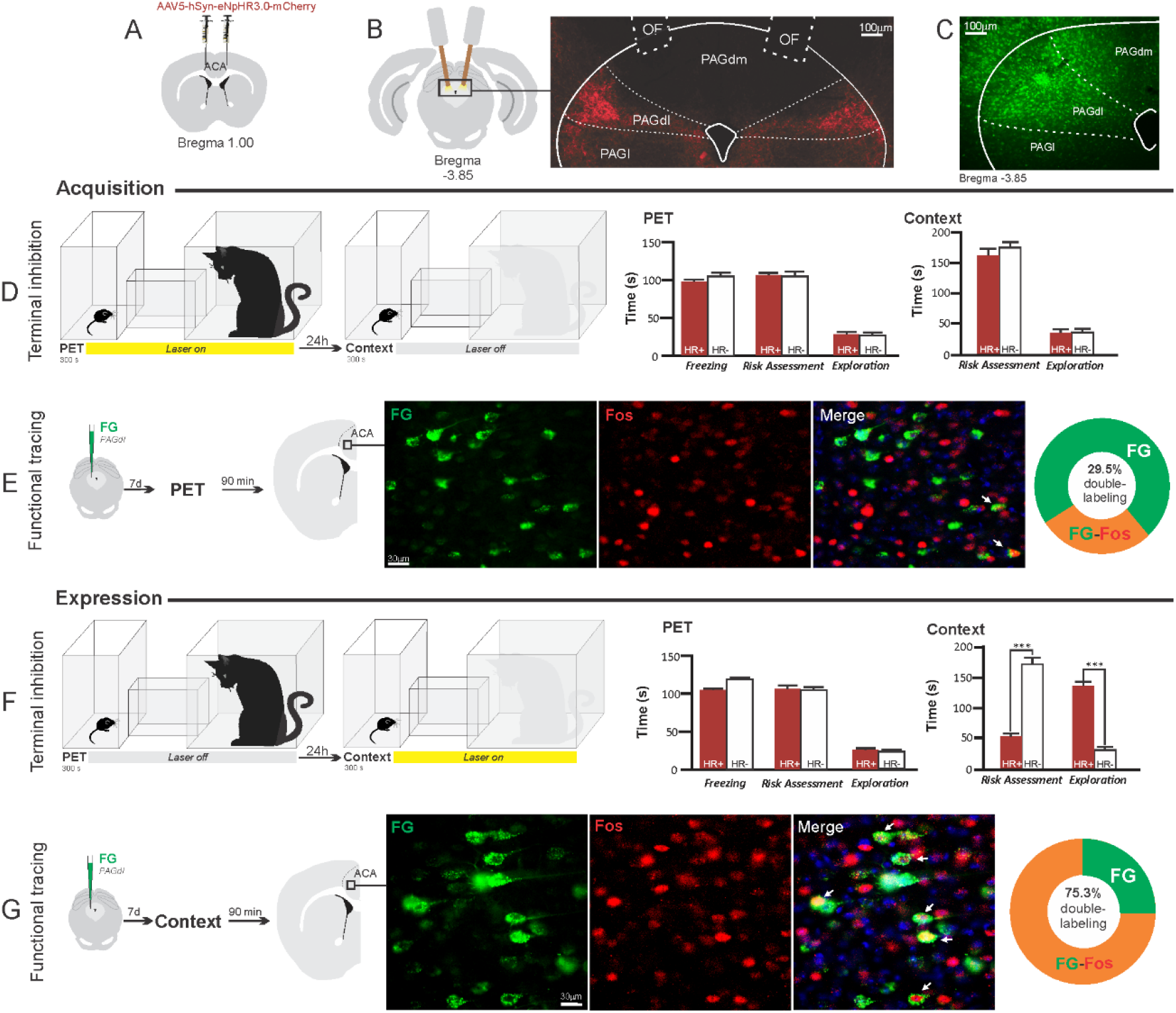
Optogenetic silencing and functional tracing of the ACA > PAGdl pathway during the acquisition and expression of contextual fear to predator threat. **A.** Schematic drawing illustrating the bilateral injection in the ACA of a viral vector expressing halorhodopsin-3.0 (eNpHR3.0) fused with mCherry. **B.** Schematic drawing (left) and fluorescence photomicrograph (right) showing the ACA projections to the PAGdl and location of bilateral optical fibers implanted close to the PAGdl (*of* - optic fibers’ tips position). **C.** Fluorescence photomicrograph illustrating the FG injection in the PAGdl for the functional tracing. **D and F.** Optogenetic silencing of the ACA > PAGdl pathway during the cat exposure (**D**) or the predatory context (**F**). Experimental design (on the left) and mean (± SEM) values of the behavioral responses during Predator Exposure (PET) and Predatory Context (on the right). For silencing during PET condition - Groups: HR+ (*n*=7) and HR-(*n*=5); for silencing during Context condition - Groups: HR+ (*n*=9) and HR-(*n*=5). Data are shown as mean ± SEM. 2 x 2 ANOVA for freezing and three-way ANOVAs for risk assessment and exploration followed by Tukey’s HSD test post hoc analysis (*p<0.001). **E and G.** Functional tracing of the ACA > PAGdl pathway during the cat exposure (acquisition phase, **E**) and the predatory context (expression phase, **G**). Left - Experimental design: unilateral FG injection in the PAGdl, and 7 days later perfusion 90 min after the cat exposure (n= 4, **E**) or the context exposure (n=4, **G**). Center - Fluorescence photomicrographs illustrating, in the ACA, FG labeled cells in green (Alexa 488), Fos protein positive cells labeled in red (Alexa 594) and merged view of the FG and FOS labeled cells (arrows indicate FG/FOS double labeled cells). Right – Graphic representation of the percentage of FG/Fos double labeled cells in the ACA. Abbreviations - ACA, anterior cingulate area; FG, Fluoro gold; of, optical fiber; PAGdl, periaqueductal gray, dorsolateral part; PAGdm, periaqueductal gray, dorsomedial part; PAGl, periaqueductal gray, lateral part; PET, predator exposure test.

We also ran functional tracing experiments. Here, each one of the ACA selected targets received a unilateral deposit of a retrograde tracer (Fluoro Gold). One week later, animals were exposed either to the cat only or to the predatory context and perfused 90 min after (Figs. 4E, G; 5E, G; 6E, G and 7E, G). For each one of these targets, we examined the percentage of Fluoro Gold labeled cells expressing Fos protein in the ACA in response to the cat exposure or exposure to the predatory context.

#### ACA > BLA projection

Our results showed that photoinhibition of the ACA > BLA projection during cat exposure did not change innate fear responses but significantly reduced contextual fear responses reducing risk assessment and increasing exploration compared to the control group (Fig 4D). Conversely, photoinhibition of the ACA > BLA projection in eNpHR3.0-expressing mice during predatory context did not change the risk assessment and exploration times compared to the control group (Fig. 4F). The results suggest that photoinhibition of the ACA > BLA projection impairs the acquisition but not the expression of contextual fear response to a predator threat. Complete statistical analysis for the behavioral data of the photoinhibition of the AM > BLA projection is supplied in the supplementary material (S7c).

Animals that received FG injection in the BLA underwent statistical analysis (chi-square test) of the functional tracing. The results revealed a significant difference between the proportion of *FG/Fos* double labeled cells in the cat exposure condition as compared to the predatory context condition (χ^2^[0.05,1]=84.12; p<0.001). The ACA contained a significantly larger percentage of double-labeled Fos/FG cells in animals exposed to the cat (83.2%) compared to those exposed to the predatory context (23.8%) (Fig. 4E, G). This suggests a larger engagement of the ACA cells projecting to the BLA during the acquisition compared to the expression of contextual fear anti-predatory responses.

#### ACA > PERI projection

The results showed that photoinhibition of the ACA > PERI projection during cat exposure did not change innate fear responses but significantly reduced contextual fear responses reducing risk assessment and increasing exploration compared to the control group (Fig 5D). Conversely, photoinhibition of the ACA > PERI projection in eNpHR3.0-expressing mice during predatory context did not change the risk assessment and exploration times versus the control group (Fig. 5F). The results suggest that silencing the ACA > PERI projection impairs the acquisition but not the expression of contextual fear response to a predator threat. Complete statistical analysis for the behavioral data of the photoinhibition of the AM > PERI projection is supplied in the supplementary material (S7d).

For the animals that received the retrograde FG injection in the PERI, the statistical analysis (chi-square test) of the functional tracing results revealed a significant difference between the proportion of *FG/Fos* double labeled cells in the cat exposure condition versus the predatory context condition (χ^2^[0.05,1]=56.552; p<0.001); here, the ACA contained a significantly larger percentage of double-labeled Fos/FG cells in animals exposed to the cat (47.3%) compared to those exposed to the predatory context (15.1%) (Fig. 5E, G). This suggests a larger engagement of the ACA cells projecting to the PERI during the acquisition compared to the expression of contextual fear responses.

#### ACA > POST projection

Compared to the control group, NpHR3.0-expressing mice that received photoinhibition of the ACA > POST projection during cat exposure or predatory context did not change the risk assessment responses and fearless exploration during exposure to the environment previously visited by a predator (Fig. 6D, F). The results suggest that silencing the ACA > POST projection apparently had no effect on the acquisition and expression of contextual fear to a predator threat. Complete statistical analysis for the behavioral data of the photoinhibition of the AM > POST projection is supplied in the supplementary material (S7e).

In line with this view, functional tracing in POST FG-injected animals revealed that the ACA contained a relatively low percentage of double-labeled Fos/FG cells in response either to the cat (21.3%) (Fig. 7E) or the predatory context (19%) (Fig. 6G). The statistical analysis (chi-square test) revealed no significant difference between the proportion of *FG/Fos* double-labeled cells in the cat exposure condition versus the predatory context condition (χ^2^[0.05,1]=0.05206; p>0.75). Thus, there is a relatively small engagement of the ACA cells projecting to the POST during both the acquisition and expression of contextual fear anti-predatory responses.

#### ACA > PAGdl projection

These results revealed that photoinhibition of the ACA > PAGdl projection during cat exposure did not change innate or contextual fear responses (Fig. 7D). Conversely, compared to the control group, photoinhibition of the ACA > PAGdl projection during exposure to the predatory context in eNpHR3.0-expressing mice yielded a significant decrease in the risk assessment and an increase in exploration (Fig. 7F). The results showed that photoinhibition of the ACA > PAGdl projection had no effect on the acquisition but impaired the expression of contextual fear to a predator threat. Complete statistical analysis for the behavioral data of the photoinhibition of the AM > PAGdl projection is supplied in the supplementary material (S7f).

For the animals that received the retrograde FG injection in the PAGdl, the statistical analysis (chi-square test) of the functional tracing results revealed a significant difference between the proportion of *FG/Fos* double labeled cells in the cat exposure condition as compared to the predatory context condition (χ^2^[0.05, 1]=79.624; p<0.001). The ACA contained a significantly larger percentage of double labeled Fos/FG cells in animals exposed to the predatory context (75.3%) (Fig. 7G) compared to those exposed to the cat (29.5%) (Fig. 7E). This suggests a larger engagement of the ACA cells projecting to the PAGdl during the expression compared to the acquisition of contextual fear responses.

## DISCUSSION

Combining the use of pharmaco- and optogenetic circuit manipulation tools and functional tracing, we untangled the ACA role in processing contextual fear memory to a predator threat.

To test the acquisition and expression of predator contextual fear responses, we used a two-phase paradigm composed of cat exposure (acquisition phase), and, on the following day, exposure to the context where the predator had been previously encountered (expression phase) (see S1 and S2). Notably, the defensive responses in mice differ somewhat from those seen in rats exposed to cat and then to the predator-associated context. During cat exposure, rats spend close to 90% of the time freezing and 2% of the time risk assessing the environment (Ribeiro-Barbosa et al., 2005); mice froze for approximately 30% of the time and risk assessed the environment for close to 30% of the time (see S1). During exposure to the predatory context, rats only froze a little and then spent time risk assessing the predator-related context for 70% of the time (Ribeiro-Barbosa et al., 2005), which is different from mice that presented no freezing and a relatively weaker risk assessment response—close to 50% of the time (see S1). An important feature of the experimental procedure used here was that the behavioral responses were very stable among the animals tested within each individual phase of the testing schedule. This feature may be, at least in part, accounted for by the long habituation period, which appears to stabilize anti-predatory behavioral responses (Ribeiro-Barbosa et al., 2005).

Pharmacogenetic inhibition of the ACA during cat exposure did not change innate responses but significantly reduced contextual fear responses. This suggests that the ACA is involved in the acquisition of predator-related contextual fear responses. In line with these results, studies using fear conditioning to physically aversive stimuli (i.e., foot shock) have also supported the idea that the ACA appears to be necessary for the acquisition of contextual fear (Tang et al., 2005; Bissière et al., 2008). Our functional tracing analysis combining retrograde tracer in the ACA and Fos immunostaining revealed that the ACA is particularly influenced by afferent sources of inputs conveying contextual information and predator cues during the acquisition phase.

Among the cortical inputs, the ventral retrosplenial (RSPv) and medial visual (VISm) areas presented the largest percentage of FG retrogradely labeled cells expressing Fos (around 50% of the retrogradely labeled cells). The retrosplenial cortex is critical for recognition of an environment directly paired with an aversive event (Robinson et al., 2018). Studies using fear conditioning to physically aversive stimuli (i.e., foot shock) have shown that the retrosplenial cortex is involved in contextual fear memory (Keene and Bucci, 2008). Recent studies using two-photon imaging revealed that landmark cues served as dominant reference points and anchored in the retrosplenial cortex the spatial code, which is the result of local integration of visual, motor, and spatial information (Fisher et al., 2020). The VISm is necessary for the visuospatial discrimination used in the integration of allocentric visuospatial cues (Sánchez et al., 1997; Espinoza et al., 1999) and may as well be involved in integrating contextual cues. Therefore, the results suggest that cortical fields projecting to the ACA particularly active during the acquisition phase of predator fear memory are involved in computing contextual landmarks.

In the thalamus, our functional tracing revealed that the central lateral (CL) and ventral anteromedial thalamic (AMv) nuclei contained more than 50% of the FG retrogradely-labeled cells expressing Fos during the acquisition phase of predator fear memory. Both the CL and AMv are likely to convey information regarding the predatory threat. The CL receives projections from the dorsal part of the periaqueductal gray (Kincheski et al., 2012), and the AMv receives substantial inputs from the dorsal premammillary nucleus (Canteras and Swanson, 1992). The dorsal premammillary nucleus is part of the predator-responsive circuit of the medial hypothalamus (Gross and Canteras, 2012) and is the most responsive site to a live predator or its odor (Cezario et al., 2008; Dielenberg et a., 2001).

In order to test whether the information relayed through the projection from the anteromedial thalamic nucleus (AM) to the ACA influences the acquisition of predator fear memory, we silenced the AM > ACA pathway during cat exposure. Photoinhibition of the AM > ACA projection during cat exposure did not change innate fear responses but significantly reduced contextual fear responses. This suggests that silencing the AM > ACA projection impairs the acquisition of contextual fear response to a predator threat. In line with this result, studies from our group have shown that pharmacological inactivation of the AMv drastically reduced the acquisition but not the expression of contextual fear responses to predator threats (de Lima et al., 2017). In contrast, anterior thalamic lesions affect contextual fear conditioning to physically aversive stimuli to a minor degree (i.e., footshock). Anterior thalamic lesions slow down the acquisition of contextual fear conditioning but do not affect contextual fear memory tested in the short term (Dupire et al, 2013; Marchand et al., 2014).

Our findings collectively support the idea that the ACA integrates contextual and predator cues during the acquisition phase to provide predictive relationships between the context and the threaten stimuli and influence memory storage. The ventral hippocampus and basolateral amygdala appear as critical sites for memory storage of contextual fear to predator threats. Ventral hippocampal cytotoxic lesions significantly reduced conditioned defensive behaviors during re-exposure to the predator-associated context (Pentkowski et al., 2006). Likewise, cytotoxic NMDA lesions in the basolateral amygdala impaired contextual fear responses to predator threat (Martinez et al., 2011; Bindi et al., 2018). In line with these ideas, recent findings from our lab indicates that cycloheximide (a protein synthesis inhibitor) injection in the basolateral amygdala or the ventral hippocampus impairs contextual fear responses to predatory threat (F. Reis and N.S. Canteras personal observations). Therefore, we investigated how the ACA would facilitate associative plasticity and memory storage in the basolateral amygdala and hippocampus. To this end, we used optogenetic silencing and functional tracing combining Fluoro Gold and Fos immunostaining to examine how the ACA entrains selective targets to influence acquisition or expression of predator fear memory.

Among the ACA targets, we start by exploring projections to the basolateral amygdala and the perirhinal region. Previous model of the prefrontal regulation of memory formation suggests that the prefrontal cortex is likely to influence associative plasticity and memory storage in the amygdala and hippocampus using two branches (Gilmartin et al., 2014). One direct branch is to the basolateral amygdala (BLA) and conveys information about the predictive value of the relevant clues during learning. The other branch is to the perirhinal cortices, which occupies a strategic position in this network influencing both the hippocampus and the amygdala. Tract-tracing studies revealed that the perirhinal region (PERI) provides dense projections to the basolateral amygdala and the hippocampal formation (Shi and Cassell, 1999). Photoinhibition of both ACA > BLA and ACA > PERI pathways during cat exposure significantly reduced contextual fear responses thus suggesting a role in the acquisition of predator fear memory. In contrast, photoinhibition of both pathways did not influence the expression of contextual fear memory. Corroborating these findings, our functional tracing revealed that both ACA > BLA and ACA > PERI paths are significantly more active during the acquisition compared to the expression of predator contextual fear memory.

The POST in turn is necessary for normal acquisition of contextual and auditory fear conditioning (Robinson and Bucci, 2012). Thus, we tested the ACA > POST pathway as a putative candidate to influence predator fear memory. However, the ACA > POST pathway's photoinhibition did not impair either the acquisition or the expression of contextual fear memory. Likewise, our functional tracing examined the percentage of Fos positive cells in the ACA projecting to the POST and revealed a relatively low percentage of double labeled cells both in the acquisition and the expression of predatory contextual fear memory. However, it is important to recall that the POST is a critical brain region to guide navigation using both visual and self-movement cues (Yoder et al., 2019). Therefore, it would be nice to investigate whether this massive ACA > POST pathway could have a role in shaping navigation in animals facing predator threats.

Pharmacogenetic inhibition of the ACA during the exposure to the predatory context significantly reduced contextual fear responses. This result further suggests that the ACA is also involved in the expression of predator-related contextual fear responses. Exposure to the predatory context yielded a different activation pattern among the sources of inputs to the ACA compared to those seen during cat exposure. Unfortunately, our functional tracing does not offer any plausible hint regarding the putative source of inputs to the ACA driving retrieval of memory traces to influence contextual fear expression. In this regard, we note that the ACA is involved in the consolidation of contextual fear conditioning to physically aversive stimuli (Einarsson and Nader, 2012). Future work could investigate whether the ACA is also involved in the consolidation of contextual fear memory to predator threat and how this consolidation would influence the retrieval predator fear memory in the ACA.

The dorsolateral periaqueductal gray (PAGdl) appears to be a likely ACA target to influence both the acquisition and expression of contextual fear responses to predator threats. Previous studies have indicated that the dorsal PAG influences the acquisition of contextual fear to predatory threat (de Andrade Rufino et al., 2019). Moreover, considering the involvement of the PAGdl in the expression of predator fear memory, it has been found that exposure to the predatory context yields a significant Fos expression in the dorsal PAG (Cezario et al., 2008)—a region where electrical, pharmacological, and optogenetic stimulation have been shown to produce freezing, flight, and risk assessment behavior in the absence of a predatory threat (Bittencourt et al., 2004; Assareh et al., 2016; Deng et al., 2016). The present findings showed that photoinhibition of the ACA > PAGdl pathway during cat exposure did not change innate or contextual fear responses. Conversely, photoinhibition of the ACA > PAGdl projection during predatory context altered contextual fear responses. In line with these findings, the functional tracing revealed that the ACA contained a significantly larger proportion of double labeled Fos/FG cells for the PAGdl FG injected animals in response to the context versus the cat exposure. Collectively, our findings revealed that the ACA > PAGdl pathway is significantly more active during exposure to the predatory context and influences the expression of contextual fear to predator threat.

Overall, the ACA can provide predictive relationships between the context and the predator threat and influences fear memory acquisition through projections to the basolateral amygdala and perirhinal region and the expression of contextual fear through projections to the dorsolateral periaqueductal gray. These findings may be applied in more general terms to understand memory processing of fear threats entraining hypothalamic circuits (i.e., such as the predator- and conspecific-responsive circuits underlying predatory and social threats, respectively) that engage the ventral anteromedial thalamic > ACA pathway (Gross and Canteras, 2019). Notably, both the predator- and conspecific-responsive hypothalamic circuits comprise the dorsal premammillary nucleus that projects densely to the ventral anteromedial thalamic nucleus (Canteras and Swanson, 1992), and therefore, engage the ventral anteromedial thalamic > ACA pathway. Of relevance, previous studies showed that the anteromedial thalamus' cytotoxic lesions impair contextual fear in a social defeat-associated context (Rangel et al., 2018). Thus, the present results open interesting perspectives for understanding how the ACA is involved in processing contextual fear memory to predator threats as well as other ethologic threatening condition such as those seen in confrontation with a conspecific aggressor during social disputes.

## METHODS

### Animals

Adult male mice, C57BL/6 (n = 192) weighing approximately 28 g were used in the present study. The mice were obtained from local breeding facilities and were kept under controlled temperature (23 C) and illumination (12-h cycle) in the animal quarters with free access to water and a standard laboratory diet. All experiments and conditions of animal housing were carried out under institutional guidelines [Colégio Brasileiro de Experimentação Animal (COBEA)] and were in accordance with the NIH Guide for the Care and Use of Laboratory Animals (NIH Publications No. 80-23, 1996). All of the experimental procedures had been previously approved by the Committee on the Care and Use of Laboratory Animals of the Institute of Biomedical Sciences, University of São Paulo, Brazil (Protocol No. 085/2012). Experiments were always planned to minimize the number of animals used and their suffering.

### Sterotaxic surgery, viral injections and optical fiber implantation

Mice were anesthetized in a box saturated with Isoforine (Cristália Laboratories, SP, Brazil) and then immediately positioned on a stereotaxic instrument (Kopf Instruments, CA, USA). The anesthesia was maintained with 1-2% Isoforine/oxygen mix and body temperature was controlled with a heating pad. Viral vectors were injected with a 5 μl Hamilton Syringe (Neuros Model 7000.5 KH). Injections were delivered at a rate of 5 nl/min using a motorized pump (Harvard Apparatus). The needle was left in place for 5 min after each injection to minimize upward flow of viral solution after raising the needle. For the photoinhibition experiments, immediately after the viral injections, optical fibers (Mono Fiber-optic Cannulae 200/230-0.48, Doric Lenses Inc. Quebeq, Canada) were implanted and fixed onto animal skulls with dental resin and micro screws (DuraLay, IL, USA). The animals were allowed 1 week to recover from the surgery, and 3 weeks later we started the behavioral experiments.

For the pharmacogenetic inhibition, ACA was injected bilaterally with 150 nl of AAV5-hSyn-HA-hM4D(Gi)-IRES-mCitrine (Dr. Bryan Roth; Addgene plasmid #50464) or AAV5-hSyn-eGFP (titer ≥ 7×10^12^ vg/mL; Addgene viral prep #50465-AAV5) as control. For the photoinhibition of the AM > ACA projection, mice were bilaterally injected into the AM (AP −0.7, ML± 0.5, DV −3.7) with 20 nl of AAV5-hSyn-eNpHR3-mCherry (titer ≥ 1×10^13^ vg/mL; University of North Carolina, Vector Core) or AAV5-hSyn-mCherry (control virus; titer ≥ 7×10^12^ vg/mL; University of North Carolina, Vector Core), and optical fibers were bilaterally implanted into the ACA (at a 10° angle from the vertical axis; AP +1.0, ML± 0.7, DV −0.6). For the photoinhibition of the ACA projections to selected targets during acquisition and expression of contextual fear responses, mice were bilaterally injected into the ACA (AP +1.0, ML ±0.3, DV −1.1) with 80 nl of AAV5-hSyn-eNpHR3-mCherry (titer ≥ 1×10^13^ vg/mL; University of North Carolina, Vector Core) or AAV5-hSyn-mCherry (control virus; titer ≥ 7×10^12^ vg/mL; University of North Carolina, Vector Core), and optical fibers were bilaterally implanted into the BLA (AP −0.95, ML±3.3, DV −3.3), PERI (AP − 3.9, ML ±0.5 from the lateral skull surface, DV −0.5 from the local brain surface), POST (at a 20° angle from the vertical axis; AP −3.9, ML ±2.7, DV −0.75) or PAGdl (at a 15° angle from the vertical axis; AP −3.85, ML ±0.9, DV −1.8). For tracing the ACA projections, mice were unilaterally injected into the ACA (AP +1.0, ML ±0.3, DV −1.1) with 80 nl of AAV5-hSyn-eNpHR3-mCherry (titer ≥ 1×10^13^ vg/mL; University of North Carolina, Vector Core).

### Experimental apparatus and behavioral procedures

The experimental apparatus was made of clear Plexigas and consisted of a Box 1 (with bedding; 15 cm long x 25 cm wide x 30 cm high) connected to a Box 2 (45 x 30 x 30 cm) by a hallway (25 x 10 x 30 cm). The Box 1 was separated from the hallway by a sliding door, and the Box 2 was divided into two compartments (15 cm long and 30 cm long) by a wall, which was removed during the predator exposure test (PET) (see Supplementary Material).

#### Habituation phase

For five days, each mouse was placed into Box 1 and left undisturbed for 5 min. Next, we opened the Box 1 sliding door, and the animal was allowed to explore the hallway and the Box 2 for 10 min. At the end of this session, the mouse was returned to its home cage.

#### Predator Exposure Test (PET)

On the 6^th^ day, a neutered 2-years-old male cat was placed and held in the Box 2 by an experimenter, and the mouse was placed into Box 1, and 5 min after, the Box 1 sliding door was opened, and the animals were exposed for 5 min to the cat. After the cat was removed at the end of the 5-min period, the hallway and the Box 2 were cleaned with 5 % alcohol and dried with paper towels, and the mouse was placed back into its home cage.

#### Context

On the day after the cat exposure, the mouse was placed back into Box 1, and 5 min after, the Box 1 sliding door was opened, and the mouse was exposed for 5 min to the environment where the predator had been previously encountered.

#### Behavior analysis

All behavioral sessions were recorded using a high-speed (120fps) camera (DMC-FZ200, Panasonic) and they were blindly scored by a trained observer using the BORIS software (Behavior Observation Research Interactive Software). The behavioral data were processed in terms of duration (total duration per session). The following behaviors were measured:

- Freezing: cessation of all movements, except for those associated with breathing;
- Risk-assessment behaviors: comprising crouch-sniff (animal immobile with the back arched, but actively sniffing and scanning the environment) and stretch postures (consisting of both stretch attend posture, during which the body is stretched forward and the animal is motionless, and stretch approach, consisting of movement directed toward the cat compartment with the animal’s body in a stretched position);
- Fearless exploration: including nondefensive locomotion and exploratory up-right position (i.e., animals actively exploring the environment, standing over the rear paws and leaning on the walls with the forepaws).

### Pharmacogenetic inhibition

Animals were previously habituated to the handling, and on the last 3 days of the habituation phase received injections of 0.2 ml of saline i.p. For the pharmacogenetic inhibition, animals were injected with 1 mg/kg of clozapine-N-oxide i.p. (CNO; Tocris Bioscience, UK) 30 min before the beginning of the behavioral test.

#### Experimental design

Animals were tested for the Acquisition and the Expression phases. For the Acquisition phase, we tested a group of animals expressing Gi-coupled hM4Di (hM4D+; n= 7) and a control group (hM4D-; n= 6), both of which were injected with CNO prior to the PET and saline prior to the Context (Figure 1B). Similarly, for the Expression phase, we tested a group of animals expressing Gi-coupled hM4Di (hM4D+; n= 7) and a control group (hM4D-; n= 5), both of which received saline injection prior to the PET and CNO injection prior to the Context exposure (Figure 1C).

### Optogenetic inhibition

Animals were previously habituated to the optogenetic cables on the last 3 days of the habituation phase of the behavioral paradigm. This procedure consisted of plugging optogenetic cables to the implanted fiber-optic cannulae and letting the animals explore the apparatus for 5 min without any additional stimulus. Optogenetic inhibition was induced by exposing animals continuously to a yellow laser (589 nm, Low-Noise DPSS Laser System, Laserglow Techonologies).

#### Experimental design

For the photoinhibition of the AM > ACA pathway, we tested a group of animals expressing halorhodopsin-3.0 (HR+; n= 8) and a control group (HR-; n= 7), both of which had the yellow laser turned ON during PET and turned OFF during the context exposure. For the photoinhibition of the ACA projections to the selected targets during acquisition and expression of contextual fear responses, animals were tested for the Acquisition and the Expression phases. For the Acquisition phase, we tested groups of animals expressing halorhodopsin-3.0 (HR+) and control groups (HR+) that had the yellow laser turned ON during PET and turned OFF during the context exposure (ACA > BLA pathway - HR+ n= 8, HR-n= 6; ACA > PERI pathway - HR+ n= 6, HR-n= 5; ACA > POST pathway - HR+ n= 6, HR-n= 5; ACA > PAGdl pathway - HR+ n= 7, HR-n= 5). Similarly, for the Expression phase, we tested groups of animals expressing halorhodopsin-3.0 (HR+) and control groups (HR+) that had the yellow laser turned OFF during PET and turned ON during the context exposure (ACA > BLA pathway - HR+ n= 8, HR-n=6); ACA > PERI pathway - HR+ n= 6, HR-n= 5; ACA > POST pathway - HR+ n= 7, HR-n=5; ACA > PAGdl pathway - HR+ n= 9, HR-n=5).

### Functional tracing

#### Fluoro Gold injection

FG injections followed the same anesthetic and stereotaxic procedures described for the viral injections. Iontophoretic deposits of 2 % solution of Fluoro Gold (FG; Fluorochrome Inc., Colo, USA) was applied unilaterally into the ACA (AP +1.0, ML +0.3, DV −1.1), BLA (AP −0.95, ML +3.3, DV −3.45), PERI (AP − 3.9, ML −0.4 from the lateral skull surface, DV −0.9 from the local brain surface), POST (at a 20° angle from the vertical axis; AP −3.9, ML +2.7, DV −1.15) or PAGdl (AP −3.85, ML +0.45, DV −2.1). Deposits were made over 5 min through a glass micropipette (tip diameter, 15 μm) by applying a +3 μA current, pulsed at 7-second intervals, with a constant-current source (Midgard Electronics, Wood Dale, Ill, USA, model CS3).

#### Experimental design

After 7 days of recovering from surgery, we started the behavioral procedures. To study the activation pattern of the different ACA source of inputs during the cat exposure and the predatory context, ACA FG injected animals were perfused 90 min after the PET (n=6) or the Context exposure (n= 6). To quantify the relative amount of activation of the ACA projections to the selected targets during the acquisition and expression of contextual fear to predator threat, FG injected animals into the selected targets were perfused 90 min after the PET (BLA n=4; PERI n=4; POST n=4; PAGdl n=4) or the Context exposure (BLA n=4; PERI n=4; POST n=4; PAGdl n=4).

### Perfusion and histological processing

After the experimental procedures, animals were deeply anesthetized in a box saturated with Isoforine and transcardially perfused with a solution of 4.0 % paraformaldehyde in 0.1 M phosphate buffer at pH 7.4. The brains were removed and left overnight in a solution of 20 % sucrose in 0.1 M phosphate buffer at 4 C. The brains were then frozen, and 5 series of 30 mm-thick sections were cut with a sliding microtome in the frontal plane. Sections from all the viruses and FG injections were taken to the fluorescent microscope to verify the injection sites and projection fields evidenced by the fluorescent reporters.

#### FG/Fos immunofluorescence double-labeling

One series of sections was washed with KPBS to remove the cryoprotectant solution and then incubated with polyclonal rabbit anti-c-Fos (PC-38; Calbiochem-Millipore) at 1:20000 in 0.5% Blocking Reagent (Roche) in KPBS for 48 h at 4 °C. Sections were then washed in KPBS and incubated with Anti-Rabbit Alexa 594 Goat IgG (H+L) (Invitrogen) at 1:500 during 2 h at random temperature (RT). Sections were then washed in KPBS and incubated with rabbit anti-FG antibody (1:5000; Chemicon International, Calif, USA) in 0.5% Blocking Reagent (Roche) in KPBS for 16 h at 4 ° C. The Sections were then washed in KPBS and incubated with Anti-Rabbit Alexa 488 Goat IgG (H+L) (Invitrogen) at 1:1000 during 1.5 h at RT. At the end, the sections were mounted in gelatin-coated slides, the nuclei were counterstained with DAPI (Sigma) at 1:20000 in TBS and cover slipped with Fluoromount (Sigma). Slides were stored at 4° C in humid chambers until image capture.

### Image capture and analysis

Brain regions of interest were determined using the *Allen Mouse Brain Atlas*. Images were captured using an epi-fluorescence microscope (NIKON, Eclipse E400) coupled to a digital camera (NIKON, DMX 1200). ImageJ public domain image processing software (FIJI v1.47f) was used for image analysis. For the documentation of the viral injection sites and projection fields of viral transfected cells, we captured only the fluorescence emitted from the fluorescent reporters. For the functional tracing studies, Fos and FG were immunodetected and labeled with Alexa 594 and Alexa 488, respectively. To determine the activation pattern of the different ACA source of inputs during the cat exposure and the predatory context, we first selected the sites containing more than 10 percent of FG labeled cells expressing Fos protein. For each one of these sites, we first delineated the borders of the selected region and FG and FG/FOS double-labeled cells were counted therein. The number of counted cells were corrected by the quantified area. To quantify the relative amount of activation of the ACA projections to the selected targets during the acquisition and expression of contextual fear to predator threat, we averaged the counting from three serial 60μm apart sections, where, in each section, we delineated the ACA borders, counted FG and FG/FOS labeled cells and corrected by the quantified area.

### Electrophysiology

#### Slice preparation

Slices were prepared using a Leica VT1000s vibratome as described previously (McKay et al., 2009; Oh et al., 2013). Briefly, mice were deeply anesthetized with isoflurane and decapitated. The brains were quickly removed, immersed in ice-cold aCSF (artificial cerebrospinal fluid) solution bubbled with 95%O_2_/5%CO_2_ and sliced. The aCSF solution containing (in mM): 125 NaCl, 2.5 KCl, 1.25 NaH_2_PO_4_, 26 NaHCO_3_, 2 CaCl_2_, 1 MgSO_4_, and 25 glucose. The cut slices were transferred immediately to a warm (37°C) submerged holding chamber filled with aCSF and kept in this chamber for 30 min. Next, the slices were allowed to return to room temperature (~25°C) for at least 40 min before electrophysiology experiments.

#### Patch-clamp

Whole cell current-clamp recordings were made in aCSF at 34±0.5°C from visualized neurons using a CMOS camera (Flash4.0, Hamamatsu) mounted on SliceScope Pro 3000 microscope (Scientifica, UK); using a long working distance 40X (0.8 NA) water-immersion objective and infrared differential interference contrast optics. Patch electrodes, yielding 4-6MΩ resistance, contained (in mM): 130 KMeSO_4_, 10 KCl, 10 HEPES, 4 ATP magnesium salt, 0.4 GTP disodium salt, with pH corrected to 7.3 with KOH and osmolarity of 295±5 mOsm. Neurons were included if they had a resting membrane potential of less than −60mV, an input resistance >25MΩ, AP amplitude of >80mV from rest (calculated from the resting potential of the cell until the peak of the AP, which was evoked by brief square pulses [2ms, 1.5nA]), and stable series resistance of <20MΩ. Resting membrane potential was measured immediately after breaking into the cell. Electrode capacitance and series resistance were monitored and compensated throughout recording; cells were held between −65 to −66mV with injected current. Data were collected using a Multiclamp 700A amplifier, pClamp 10.7 software and digitized (10kHz) using a Digidata 1440 AD converter. All from Molecular Devices (USA). Data were analyzed using Clampfit 10.7 software (Molecular Devices, USA). Input resistance was calculated as the slope of the V-I curve using 500ms current steps from ⫺300pA to 0pA at 50pA steps. Action potentials (APs) were evoked using ramp current pulses (100ms pulses ranging from 100 to 500pA at 100pA steps).

#### Optogenetics

The transfected neurons with the AAV5-hSyn-eNpHR3-mCherry or AAV5-hSyn-mCherry were identified and stimulated using a pE-2 light source (CoolLED, UK). The wavelength of 585nm was used to identify and activate halorhodopsin-transfected cells. The wavelength was filtered using a double-band fluorescence cube (59022-filter set; Chroma, USA).

#### Pharmacogenetic

The neurons transfected with the pAAV-hSyn-HA-hM4D(Gi)-IRES-mCitrine or AAV5-hSyn-eGFP were identified using a pE-2 light source (CoolLED, UK). The wavelength of 470nm, filtered by the 59022 cube-set (Chroma, USA) was used to identify the transfected cells in the brain slices. CNO was superfused at a flow rate of ~3ml/min.

### Statistical analysis for Fos-FG double-immunofluorescence

To evaluate the potential discrepancy between the proportion of *FG/Fos* double labeled cells in the cat exposure condition in comparison to the predatory context condition, we employed a 2×2 chi-square test with Yates’ correction for continuity, separately for each injection site. To maintain the overall type I error at 5%, the significance level employed in each test was adjusted downward (Bonferroni’s correction) according to the total number of analyses performed (α = 0.0125). The 95% confidence intervals for proportions were calculated using ESCI – Exploratory Software for Confidence Intervals (Cumming, 2012).

### Statistical analysis for behavioral measurements

After testing for homogeneity of variance (Levine’s test), the behavioral data were square-root transformed whenever the null hypothesis of homoscedasticity was rejected. For all experiments, the analysis was performed by means of a parametric univariate analysis of variance (ANOVA), followed by a post hoc analysis (Tukey’s HSD test) when appropriate. The specific design varied depending on the structure of each experiment, ranging from one-way to three-way ANOVAs. Due to an expressive number of analyses performed and to maintain the overall type I error at 5%, the significance level employed in each ANOVA was adjusted downward by means of a Bonferroni’s correction (α = 0.0028). The average results are expressed as the mean ± SEM throughout the text and effect sizes are expressed as partial eta-squared. Two-tailed tests were used throughout the statistical analyses for both cell counting and behavioral measurements.

## Acknowledgments

This research was supported by Fundação de Amparo à Pesquisa do Estado de São Paulo (FAPESP) Research Grants #2014/05432-9 (to NSC) and #2019/27245-0 (to FAO). FAPESP fellowships to MAXL (#2016/10389-0).

## Author Contributions

MAXL and NSC - conceptualized the project, designed the study, analyzed data, interpreted results prepared and edited the manuscript. FAO performed and analyzed the patch clamp studies and MVCB analyzed data and performed the statistical analysis.

## Financial Disclosures/Conflict of Interest

The authors have no competing financial interests or potential conflicts of interest.

## Notes

### Competing Interest Statement

The authors have declared no competing interest.

