## Supplementary Material for "The anterior cingulate cortex and its role in controlling contextual fear memory to predatory threats"

### Supplementary data and material

**S1. Exposure to the predator and predatory context.** In the present study, mice were exposed to a predator (a live cat) and tested for innate and contextual fear responses. Animals were tested in an experimental apparatus that consisted of a 25 × 15 × 30 cm home cage (Box1) connected to another 30 × 45 × 30 cm chamber (Box 2) by a hallway that was 10 cm wide, 25 cm long, and 30 cm high (Fig. 1). As shown in Fig. 2A, during 5 days before testing, animals were habituated to the apparatus. The relatively long habituation period seems necessary to optimally stabilize both innate and contextual defensive responses [1]. On the 6$^{th}$ day, a neutered 2-years-old male cat was placed and held in the Box 2 by an experimenter, and the mouse was placed into Box 1, and 5 min after, the Box 1 sliding door was opened, and the animals were exposed for 5 min to the cat (Fig. 2B, Predator Exposure Test – PET Condition). Animals exposed to the cat presented clear innate defensive responses, they froze for approximately 30 percent of the time and presented clear risk assessment responses, characterized by crouch sniff and stretch postures toward the cat, close to 30 percent of the time (Table 1). After the cat was removed at the end of the 5-min period, the hallway and the Box 2 were cleaned with 5 % alcohol and dried with paper towels, and the mouse was placed back into its home cage. On the following day, animals were exposed to the same environment of the cat exposure during 5 min (Fig. 2C, Context Condition) and, presented risk assessment responses close to 50 percent of the observational period, but no freezing. Similar to previous results [1], contextual fear responses to a predator threat are characterized mostly by risk assessment behaviors. In the case of mice, we observed only risk assessment with no freezing, and in rats exposed to the predatory context, it has been reported a great deal of risk assessment with a reduced amount of freezing [1].

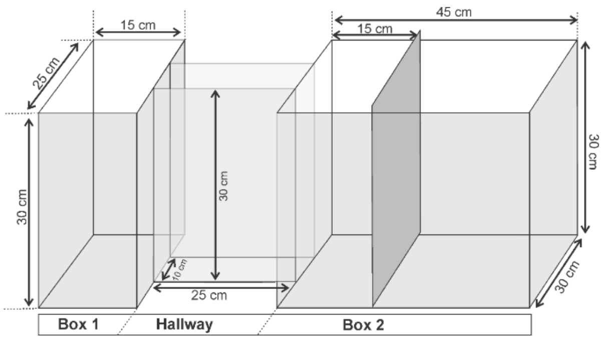


**Supplementary Figure 1. Experimental apparatus used for exposure to the Predator and Predatory Context**

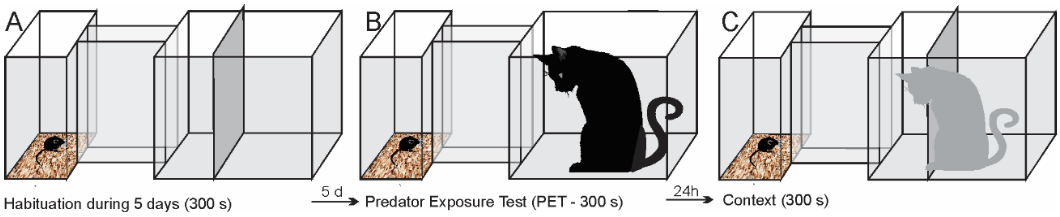


**Supplementary Figure 2. Timeline to illustrate the experimental procedure.**

**Supplementary Table 1. Behavioral data from a control group (n= 6) during the PET and Context conditions.**

| | **PET** | **Context** |
|---|---|---|
| ***Freezing*** | 102.8 ± 4.06 | 0.0 ± 0.0 |
| ***Risk assessment*** | 107.6 ± 3.52 | 161.6 ± 8.28 |
| ***Exploration*** | 25.77 ± 2.46 | 37.93 ± 4.44 |

**S2. Video - Exposure to the predator and predatory context.** Video to illustrate the behavioral responses of a male mice (C57BL/6) exposed to the cat (PET condition) and the predator-associated environment (Context condition).

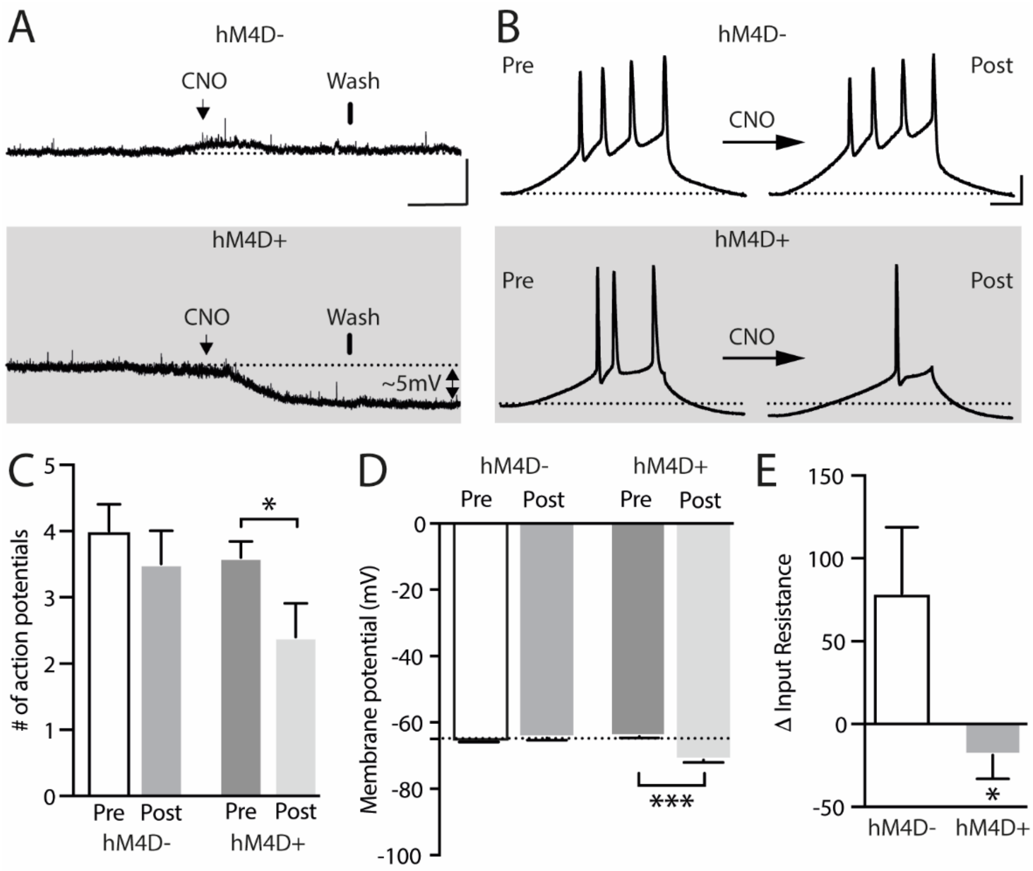


**Supplementary Figure 3. Impact of CNO on resting membrane potential and neuronal excitability in hM4D+ transfected neurons. A**: Representative traces of resting membrane potential from hM4D- neurons (control - upper panel) or hM4D+ neurons (lower panel). hM4D+ neurons hyperpolarized as they underwent extracellular CNO. The arrows show the application of the CNO (lasting around 3 minutes) and the bars show the start of drug washing. Scale bar for panel A: 10mV, 1.5min. **B**: A ramp protocol (100ms pulse of 500pA) was used to assess cellular excitability after applying CNO. Representative traces of APs evoked during the ramp protocol in hM4D- and hM4D+ neurons before and after CNO application. **C**: Bar graph shows a significant decrease in the triggering of APs after 10 µM CNO in hM4D+ neurons (pre: 3.6±0.2 and post 2.4±0.5; n= 5) when compared to hM4D- neurons (pre: 4±0.4 and post 3.5±0.5; n= 5). **D and E**: Quantification of the reduction in the resting membrane potential (hM4D-: pre: -65.8±0.14 and post: -64.7±0.6; n= 5; hM4D+: pre: -66.1±0.4 and post: -71.3±0.8; n= 6; ***p<0.001, paired t-test) and input resistance (hM4D-: 78.8±40; n= 5; hM4D+: -18.8±14.2; n= 5; *p<0.05, unpaired t-test) caused by the application of CNO. The input resistance was calculated as the slope of the I-V curve and quantified before and after the application of CNO, the difference ($\Delta = R_{in}final - R_{in}initial$) being reported in the graph. Bars represent mean±SEM. The dotted line on panels A, B and D represents -65mV.

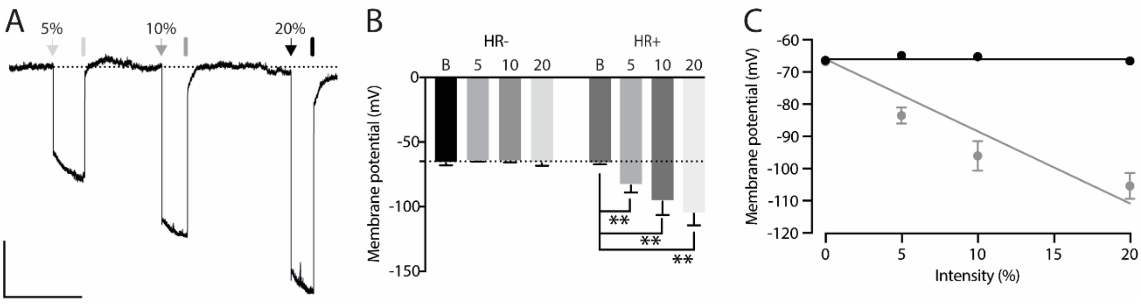


**Supplementary Figure 4. Light-induced hyperpolarization in Halorhodopsin positive neurons (HR+). A**: Representative trace of membrane potential from a HR+ neuron. The arrows and bars represent the light on and off, respectively. The percentage on the top of the panel indicates the intensity of the light which correspond to 12.8 (5%), 25 (10%) and 50.6 (20%)µW. Scale bar for panel A: 10mV, 0.5min. **B**: Membrane potential recorded during 585nm-lights on in the control (HR-) (Basal: -66.4±0.8; 5%: -64.9±0.2; 10%: -65.27±0.3 and 20%: -66.6±0.8; n= 5) and HR+ neurons (Basal: -66.8±0.2; 5%: -83.5±2.5; 10%: -96.1±4.6 and 20%: -105.4±4; n= 5, **p< 0.01). Hyperpolarization was quantified in the last 100ms before turning off the light. Bars represent mean±SEM. Dotted line represents -65mV. **C**: Linear regression of hyperpolarization caused by halorhodopsin activation due to different light intensities (HR-: slope= -0.001; HR+: slope= -2.241. Using a holding potential of -66mV as a constraint). Points represent mean±SEM.

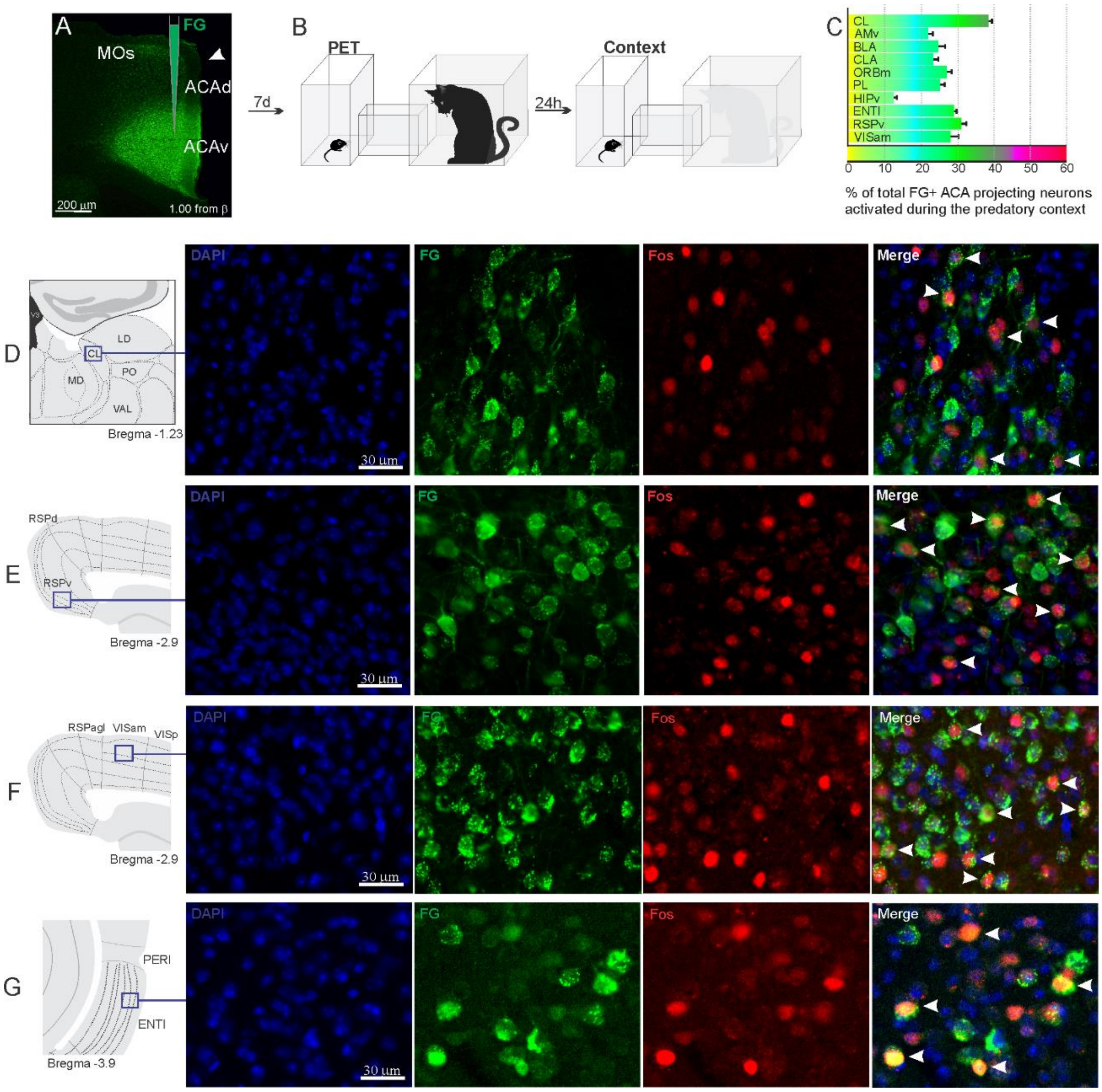


**Supplementary Figure 5. Pattern of activation of the different sources of inputs to the ACA during the exposure to predatory context.** Animals received unilateral deposit of a retrograde tracer (Fluoro Gold) in the ACA (n=6, **A**). Seven days later, animals were exposed to the cat (PET) and 24 hs later to the Predatory Context (**B**) and perfused 90 min after. **C.** Bar chart presents, for each designated structure, the proportion of activated neurons (*FG/Fos* double labeled cells) during the Context condition, among the total of FG retrogradely labeled cells (error bars indicate 95% confidence interval for a proportion). **D-G.** Schematic drawings from *Allen Mouse Brain Atlas* to show the sites containing the largest proportion of *FG/Fos* double labeled cells, followed by fluorescence photomicrographs illustrating DAPI-staining, FG labeled cells in green (Alexa 488), Fos protein positive cells labeled in red (Alexa 594) and merged view of the FG and FOS labeled cells, where arrow heads indicate FG/FOS double labeled cells. Abbreviations - ACAd, anterior cingulate area, dorsal part; ACAv, anterior cingulate area, ventral part; AMv, anteromedial thalamic nucleus, ventral part; BLA, basolateral amygdalar nucleus; CL, central lateral nucleus of the thalamus; CLA, claustrum; ENTl, entorhinal area, lateral part; FG, Fluoro gold; HIPv, hippocampus, ventral part; LD, lateral dorsal nucleus of the thalamus; MD, mediodorsal nucleus of the thalamus; MOs, secondary motor area; ORBm, orbital area, medial part; PET, predator exposure test; PERI, perirhinal area; PO, posterior complex of the thalamus; PL, prelimbic area; RSPagl, retrosplenial área, lateral agranular part; RSPd, retrosplenial area, dorsal part; RSPv, retrosplenial area, ventral part; VAL, ventral anterior-lateral complex of the thalamus; VISam, anteromedial visual area; VISp, primary visual area.

**S6. ACA projections.** Here, we provide an analysis of the ACA projections to the anteromedial visual and retrosplenial areas, as well as the sites presently investigated as putatively involved in the acquisition and/or expression of contextual fear responses to predatory threats, namely the basolateral amygdalar nucleus (BLA), the perirhinal area (PERI), the postsubiculum (POST) and the dorsolateral part of the periaqueductal grey (PAGdl). To this end, mice (n=3) were unilaterally injected into the ACA (AP +1.0, ML ±0.3, DV -1.1) with 80 nl of AAV5-hSyn-eNpHR3-mCherry. All injections were largely confined to the ACA including its dorsal and ventral parts at the level the receive the densest projection from the ventral part of the anteromedial thalamic nucleus (AMv). All cases revealed similar pattern of projections along the brain, and we chose the one with the densest terminal fields.

ACA projections to the anteromedial visual and retrosplenial areas target mostly layers I and III (Fig.). In the basolateral amygdalar complex, the ACA projects densely to the basolateral nucleus, and avoids the lateral nucleus. In the perirhinal area, a dense projection was found in the in the middle layers (II-V), spreading ventrally, to a lesser degree, to the lateral entorhinal area. In the postsubiculum, the ACA provides a dense bilaminar pattern of projection aimed at the superficial and deep layers. In the periaqueductal gray, the ACA provides particularly dense projection fields to the dorsolateral part, extending, to a lesser degree, to the lateral and dorsomedial parts. Notably, retrograde findings revealed that retrogradely Fluoro Gold labeled cells were mostly found in the ACA supragranular layers in FG deposits made in the BLA, PERI and POST, whereas the ACA cells projecting to the PAGdl were mostly found in the infragranular layers.

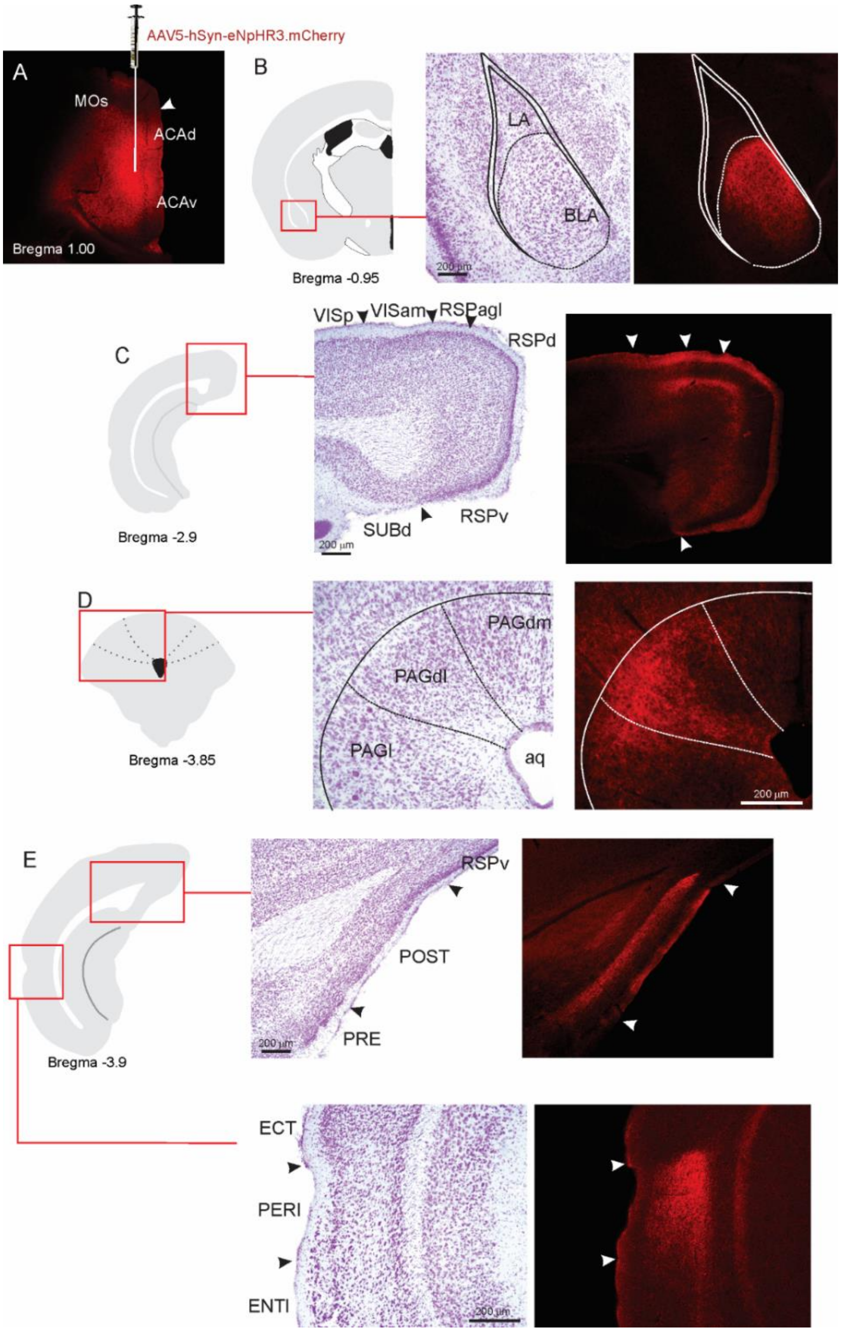


**Supplementary Figure 6. ACA projections. A.** Fluorescence photomicrograph illustrating the extent and location of unilateral viral injection in the ACA. **B – E.** On the left, schematic drawings from the *Allen Mouse Brain Atlas* to indicate the locations of the higher magnification nissl- stained and accompanying fluorescence photomicrographs (on the right) to show the ACA projections to the BLA (**B**), anteromedial visual and retrosplenial areas (**C**), PAGdl (**D**), and POST and PERI (**E**). Abbreviations: ACAd, anterior cingulate area, dorsal part; ACAv, anterior cingulate area, ventral part; Aq, aqueduct; BLA, basolateral amygdalar nucleus; ECT, ectorhinal area; ENTl, entorhinal area, lateral part; LA, lateral amygdalar nucleus; MOs, secondary motor area; PAGdl, periaqueductal gray, dorsolateral part; PAGdm, periaqueductal gray, dorsomedial part; PAGl, periaqueductal gray, lateral part; PERI, perirhinal area; POST, postsubiculum;

PRE, presubiculum; RSPagl, retrosplenial área, lateral agranular part; RSPd, retrosplenial area, dorsal part; RSPv, retrosplenial area, ventral part; SUBd, subiculum, dorsal part; VISam, anteromedial visual area; VISp, primary visual area.

**S7. Statistical analysis for behavioral experiments.**

***S7a. Pharmacogenetic inhibition of the ACA during cat exposure or predatory context.*** For the freezing, a 2x2 ANOVA revealed neither main effects for the factors virus (hM4D+ and hM4D-, $F[1,21]=1.67$; $p=0.21$; $\eta^2_p=0.073$) and phase of treatment (CNO/PET and CNO/Context, $F[1,21]=1.12$; $p=0.30$; $\eta^2_p=0.051$) nor an interaction between them ($F[1,21]=0.064$; $p=0.803$; $\eta^2_p=0.003$). For the risk assessment, a three-way ANOVA revealed a significant main effect for the factor virus (hM4D+ and hM4D-, $F[1,21]=200.75$; $p<0.001$; $\eta^2_p=0.905$), no main effect for the factors phase of treatment (CNO/PET and CNO/Context, $F[1,21]=1.01$; $p=0.32$; $\eta^2_p=0.046$) and exposure (PET and Context, $F[1,21]=3.47$; $p=0.07$; $\eta^2_p=0.142$), and a significant interaction between the factors virus and exposure ($F[1,21]=144.5$; $p<0.001$; $\eta^2_p=0.873$). Post hoc pairwise comparisons (Tukey's HSD test) for the animals tested during the predatory context revealed for the hM4D+ animals that received CNO during cat exposure (Figure 1B) and the hM4D+ group that received CNO during the predatory context (Figure 1C) a significant decrease in risk assessment compared to the control hM4D- group ($p<0.001$). For the relaxed exploration, a three-way ANOVA revealed a significant main effect for the factors virus (hM4D+ and hM4D-, $F[1,21]= 178.74$; $p<0.001$; $\eta^2_p=0.895$) and exposure (PET and Context, $F[1,21]=429.58$; $p<0.001$; $\eta^2_p=0.953$), but no main effect for the factor phase of treatment (CNO/PET and CNO/Context, $F[1,21]= 1.41$; $p= 0.25$; $\eta^2_p=0.063$), and a significant interaction between the factors virus and exposure ($F[1,21]= 188.22$; $p<0.001$; $\eta^2_p=0.899$). Post hoc pairwise comparisons (Tukey's HSD test) for the animals tested during the predatory context revealed for the hM4D+ animals that received CNO during cat exposure (Figure 1B) and the hM4D+ group that received CNO during the predatory context (Figure 1C) a significant increase in relaxed exploration compared to the control hM4D- group ($p<0.001$).

***S7b. Photoinhibition of the AM > ACA projection during cat exposure***. For the freezing, a one-way ANOVA revealed no main effect for the factor virus (HR+ and HR-, $F[1,13]=$

1.05; p= 0.324; $\eta^2_p$=0.074). For the risk assessment, a 2x2 ANOVA revealed a main effect for the factor virus (HR+ and HR-, F[1,13]=149.79; p<0.001; $\eta^2_p$=0.920) and a significant interaction between the factors virus and exposure (F[1,13]=178.43; p<0.001; $\eta^2_p$=0.932). Post hoc pairwise comparisons (Tukey's HSD test) revealed for the HR+ animals a significant decrease in the risk assessment during the context exposure (p<0.001) (Figure 4E). For exploration, a 2x2 ANOVA revealed a main effect for the factor virus (HR+ and HR-, F[1,13]=102.18; p<0.001; $\eta^2_p$=0.887) and a significant interaction between the factors virus and exposure (F[1,13]=238.00; p<0.001; $\eta^2_p$=0.948). Post hoc pairwise comparisons (Tukey's HSD test) revealed for the HR+ animals a significant increase in the exploration during the context exposure (p<0.001) (Figure 4E).

***S7c. Photoinhibition of the ACA > BLA projection during cat exposure or predatory context***. For the freezing, a 2x2 ANOVA revealed neither main effects for the factors virus (HR+ and HR-, F[1,24]=2.98; p=0.097; $\eta^2_p$=0.110) and phase of treatment (Photoinhibition /PET and Photoinhibition /Context, F[1,24]=0.20; p=0.656; $\eta^2_p$=0.008) nor an interaction between them (F[1,24]=0.053; p=0.819; $\eta^2_p$=0.002). For the risk assessment, a three-way ANOVA revealed significant main effects for the factors virus (HR+ and HR-, F[1,24]=78.6; p<0.001; $\eta^2_p$=0.766), phase of treatment (Photoinhibition/PET and Photoinhibition/Context, F[1,24]=23.83; p<0.001; $\eta^2_p$=0.498) and exposure (PET and Context, F[1,24]=35.38; p<0.001; $\eta^2_p$=0.596), as well as a significant three-way interaction among these factors (F[1,24]=45.74; p<0.001; $\eta^2_p$=0.656). For the animals tested during the predatory context, post hoc pairwise comparisons (Tukey's HSD test) revealed for the HR+ animals that received photoinhibition during cat exposure a significant decrease in risk assessment compared to the control HR- group (p<0.001) (Fig. 5D), whereas the HR+ group that received photoinhibition during the context did not differ from the control HR- group (p=0.99) (Fig. 5F). For the exploration, a three-way ANOVA revealed significant main effects for the factors virus (HR+ and HR-, F[1,24]=106.76; p<0.001; $\eta^2_p$=0.816), phase of treatment (Photoinhibition/PET and Photoinhibition/Context, F[1,24]=70.91; p<0.001; $\eta^2_p$=0.747) and exposure (PET and Context, F[1,24]=256.10; p<0.001; $\eta^2_p$=0.914), as well as a significant three-way interaction among these factors (F[1,24]=93.76; p<0.001; $\eta^2_p$=0.796). For the animals tested during the predatory context, post hoc pairwise comparisons (Tukey's HSD test) revealed for the HR+ animals that received photoinhibition during cat exposure a significant increase in the relaxed exploration compared to the control HR- group ($p<0.001$) (Fig. 5D), whereas the HR+ group that received photoinhibition during the context did not differ from the control HR- group ($p=0.99$) (Fig. 5F).

***S7d. Photoinhibition of the ACA > PERI projection during cat exposure or predatory context.*** For the freezing, a 2x2 ANOVA revealed neither main effect for the factor virus (HR+ and HR-, $F[1,18]=0.259$; $p=0.617$; $\eta^2_p=0.014$) and the factor phase of treatment (Photoinhibition /PET and Photoinhibition /Context, $F[1,18]=4.46$; $p=0.049$; $\eta^2_p=0.198$), and no significant interaction between them ($F[1,18]=3.24$; $p=0.088$; $\eta^2_p=0.152$). For the risk assessment, a three-way ANOVA revealed significant main effects for the factors virus (HR+ and HR-, $F[1,18]=61.56$; $p<0.001$; $\eta^2_p=0.773$) and phase of treatment (Photoinhibition/PET and Photoinhibition/Context, $F[1,18]=63.25$; $p<0.001$; $\eta^2_p=0.778$), but not for the factor exposure (PET and Context, $F[1,18]=7.69$; $p=0.0125$; $\eta^2_p=0.299$). There was a significant three-way interaction among these factors ($F[1,18]=19.36$; $p<0.001$; $\eta^2_p=0.518$). For the animals tested during the predatory context, post hoc pairwise comparisons (Tukey's HSD test) revealed for the HR+ animals that received photoinhibition during cat exposure a significant decrease in risk assessment compared to the control HR- group ($p<0.001$) (Fig. 6D), whereas the HR+ group that received photoinhibition during the context did not differ from the control HR- group ($p=0.96$) (Fig. 6F). For the exploration, a three-way ANOVA revealed significant main effects for the factors virus (HR+ and HR-, $F[1,18]=135.44$; $p<0.001$; $\eta^2_p=0.883$), phase of treatment (Photoinhibition/PET and Photoinhibition/Context, $F[1,18]=204.85$; $p<0.001$; $\eta^2_p=0.919$) and exposure (PET and Context, $F[1,18]=343.98$; $p<0.001$; $\eta^2_p=0.950$), as well as a significant three-way interaction among these factors ($F[1,18]=127.74$; $p<0.001$; $\eta^2_p=0.876$). For the animals tested during the predatory context, post hoc pairwise comparisons (Tukey's HSD test) revealed for the HR+ animals that received photoinhibition during cat exposure a significant increase in the relaxed exploration compared to the control HR- group ($p<0.001$) (Fig. 6D), whereas the HR+ group that received photoinhibition during the context did not differ from the control HR- group ($p=0.99$) (Fig.6F).

***S7e. Photoinhibition of the ACA > POST projection during cat exposure or predatory context.*** For the freezing, a 2x2 ANOVA revealed neither main effects for the factors virus (HR+ and HR-, F[1,19]=1.08; p=0.311; $\eta^2_p$=0.054) and phase of treatment (Photoinhibition /PET and Photoinhibition /Context, F[1,19]<0.001; p=0.997; $\eta^2_p$<0.001) nor an interaction between them (F[1,19]=0.85; p=0.367; $\eta^2_p$=0.043). For the risk assessment, a three-way ANOVA revealed no main effects for the factors virus (HR+ and HR-, F[1,19]=0.37; p=0.551; $\eta^2_p$=0.019) and phase of treatment (Photoinhibition/PET and Photoinhibition/Context, F[1,19]=1.64; p=0.215; $\eta^2_p$=0.079), and a significant main effect for the factor exposure (PET and Context, F[1,19]=39.56; p<0.001; $\eta^2_p$=0.675). There was no significant three-way interaction among these factors (F[1,19]=0.018; p=0.894; $\eta^2_p$<0.001). Post hoc pairwise comparisons (Tukey's HSD test) for the animals tested during the predatory context revealed no difference for the HR+ and HR- animals that received photoinhibition during cat exposure (p=0.994) or during the context (p=1) (Fig. 7D, F). For the exploration, a three-way ANOVA revealed no significant main effects for the factors virus (HR+ and HR-, F[1,19]=2.63; p=0.121; $\eta^2_p$=0.122) and phase of treatment (Photoinhibition/PET and Photoinhibition/Context, F[1,19]=0.19; p=0.663; $\eta^2_p$=0.10), and a significant main effect for the factor exposure (PET and Context, F[1,19]=18.18; p<0.001; $\eta^2_p$=0.489). There was no significant three-way interaction among these factors (F[1,19]=1.21; p=0.285; $\eta^2_p$=0.059). Post hoc pairwise comparisons (Tukey's HSD test) for the animals tested during the predatory context revealed no difference for the HR+ and HR- animals that received photoinhibition during cat exposure (p=0.728) or during the context (p=0.999) (Fig. 7D, F).

***S7f. Photoinhibition of the* ACA > PAGdl *projection during cat exposure or predatory context.*** For the freezing, a 2x2 ANOVA revealed neither main effects for the factors virus (HR+ and HR-, F[1,22]=1.40; p=0.249; $\eta^2_p$=0.059) and phase of treatment (Photoinhibition /PET and Photoinhibition /Context, F[1,22]=1.11; p=0.302; $\eta^2_p$=0.048) nor an interaction between them (F[1,22]=0.01; p=0.923; $\eta^2_p$<0.001). For the risk assessment, a three-way ANOVA revealed significant main effects for the factors virus (HR+ and HR-, F[1,22]=38.05; p<0.001; $\eta^2_p$=0.634), phase of treatment (Photoinhibition/PET and Photoinhibition/Context, F[1,22]=33.25; p<0.001; $\eta^2_p$=0.602) and exposure (PET and Context, F[1,22]=77.22; p<0.001; $\eta^2_p$=0.778), as well as a significant three-way interaction among these factors (F[1,22]=45.23; p<0.001; $\eta^2_p$=0.673). For the animals tested during the predatory context, post hoc pairwise comparisons (Tukey's HSD test) revealed for the HR+ animals that received photoinhibition during the context a significant decrease in risk assessment compared to the control HR- group ($p<0.001$) (Fig. 8F), whereas the HR+ group that received photoinhibition during cat exposure did not differ from the control HR- group ($p=0.852$) (Fig. 8D). For the exploration, a three-way ANOVA revealed significant main effects for the factors virus (HR+ and HR-, $F[1,22]=90.20$; $p<0.001$; $\eta^2_p=0.804$), phase of treatment (Photoinhibition/PET and Photoinhibition/Context, $F[1,22]=64.93$; $p<0.001$; $\eta^2_p=0.747$) and exposure (PET and Context, $F[1,22]=103.45$; $p<0.001$; $\eta^2_p=0.825$), as well as a significant three-way interaction among these factors ($F[1,22]=62.25$; $p<0.001$; $\eta^2_p=0.739$). For the animals tested during the predatory context, post hoc pairwise comparisons (Tukey's HSD test) revealed for the HR+ animals that received photoinhibition during the context a significant increase in the exploration compared to the control HR- group ($p<0.001$) (Fig. 8F), whereas the HR+ group that received photoinhibition during the cat exposure did not differ from the control HR- group ($p=0.99$) (Fig. 8D).
